# Robust range of auditory periphery development, eye opening, and brain gene expression in Wistar rat pups that experience variation in maternal backgrounds

**DOI:** 10.1101/2020.02.17.953117

**Authors:** Jingyun Qiu, Preethi Singh, Geng Pan, Annalisa de Paolis, Frances A. Champagne, Jia Liu, Luis Cardoso, Adrián Rodríguez-Contreras

## Abstract

The experience of variation in maternal licking and grooming (LG) is considered a critical influence in neurodevelopment related to stress and cognition, but little is known about its relationship to early sensory development. In this study, we used a maternal selection approach to test the hypothesis that differences in LG during the first week of life influence the timing of hearing onset in Wistar rat pups. We performed a range of tests, including auditory brainstem responses (ABR), tracking of eye opening (EO), micro-CT X-ray tomography, and qRT-PCR to monitor neurodevelopmental changes in the female and male progeny of low-LG and high-LG dams. Our results show that variation in maternal LG is not overtly associated with different timing of ABR onset and EO in the progeny. However, the data provide insight on the delay between hearing onset and EO, on key functional and structural properties that define hearing onset at the auditory periphery, and on changes in brain gene expression that include the first evidence that: a) the hypoxia-sensitive pathway is regulated in subcortical and cortical auditory brain regions before hearing onset, and b) implicates maternal LG in regulation of Bdnf signaling in auditory cortex after hearing onset. Altogether, these findings provide a baseline to evaluate how factors that severely disrupt the early maternal environment may affect the expression of robust developmental sensory programs.

**SIGNIFICANCE STATEMENT:** Early life experience during sensitive developmental periods can induce long-term effects on the neurobiological development of the offspring. In the present work we tested the hypothesis that variation in maternal licking and grooming (LG) affects the timing of hearing onset in Wistar rat pups. To our surprise the results did not support the hypothesis. Instead, we found a robust range of early and late auditory development that was independent of maternal LG. Nevertheless, the study provides new findings on the delay between hearing onset and eye opening, on key functional and structural properties that define hearing onset at the auditory periphery, and the first evidence that a) the hypoxia-sensitive pathway is regulated in the central auditory system during the sensitive period before hearing onset, and b) maternal LG is implicated in regulation of Bdnf signaling during the sensitive period after hearing onset. These findings provide a baseline to evaluate how factors that severely disrupt the early maternal environment may affect the expression of robust developmental sensory programs.

## INTRODUCTION

In several mammalian species, including humans, maternal care is the main source of nutritional, social, and sensory stimulation that is important for survival and has the potential to impact the neurobiological development of the offspring (Curley and Champagne, 2016; González-Mariscal and Melo, 2017). Variation in rat postpartum maternal licking and grooming (LG) has been used as a model to select dams with individual differences in LG behavior and study the developmental re-programming of the offspring’s adult stress response (Liu et al., 1997; Francis et al., 1999; Weaver et al., 2004; Hancock et al., 2005; Barha et al., 2007; Menard and Hakvoort, 2007; Parent and Meaney, 2008; Walker et al., 2008; Cameron et al., 2008; Sakhai et al., 2011). However, characterization of the effects of the rearing experiences provided by dams with different LG phenotypes is incomplete, particularly with respect to how maternal LG may affect sensory development of the offspring during sensitive periods of early postnatal development, when various environmental challenges can severely disrupt mother infant interactions, reduce the chances of survival, and cause severe long-term neurobiological deficits in the progeny (Salmaso et al., 2014; Careaga, Murai and Bauman, 2017).

In a previous study, Adise *et al*. (2014) showed that a 15-minute separation followed by return to the biological or a foster mother accelerated auditory periphery development in Wistar rat pups. The effects were stronger when pups were manipulated at postnatal day 5 (P5), and weaker when pups were manipulated at P1 or P9, suggesting that the effects of maternal separation on auditory development were restricted to a sensitive period of postnatal development that occurs one week before the onset of hearing in this species. However, the contributing factors from the maternal environment, such as specific changes in maternal behavior or physiology were not identified.

In the present study, we tested the hypothesis that variation in maternal LG during the first week of life is associated with differences in the timing of hearing onset in Wistar rat pups. This idea is motivated by previous findings that maternal LG is increased in adoptive Wistar rat dams (Maccari et al., 1995), and that massage treatment during the sensitive period before eye opening (EO) accelerates development of visually evoked potentials in Long-Evans rats (Guzzeta et al., 2009). If the frequency of maternal LG influences the timing of hearing onset in the offspring, then pups within a given litter would have an early or late hearing onset that correlates with the LG phenotype of their mother (Figure 1A). To determine the relationship between maternal LG and neurodevelopmental changes in the progeny, we performed tests of auditory brainstem response (ABR), tracking of eye opening (EO), imaging development of the middle ear cavity using micro-CT X-ray tomography, and monitored changes in gene expression in auditory brainstem and primary sensory cortex of pups reared by low-LG or high-LG dams (Figure 1B-D). Contrary to our expectations, the results show that variation in maternal LG is not overtly associated with different timing of ABR onset and EO in the progeny. Nevertheless, the data provide insight on the delay between hearing onset and EO, on key functional and structural properties that define hearing onset at the auditory periphery, and the first evidence that genes of the hypoxia-sensitive pathway and Bdnf signaling are regulated during sensitive periods that occur before and after hearing onset, respectively, in Wistar rat pups.

**Figure 1.**
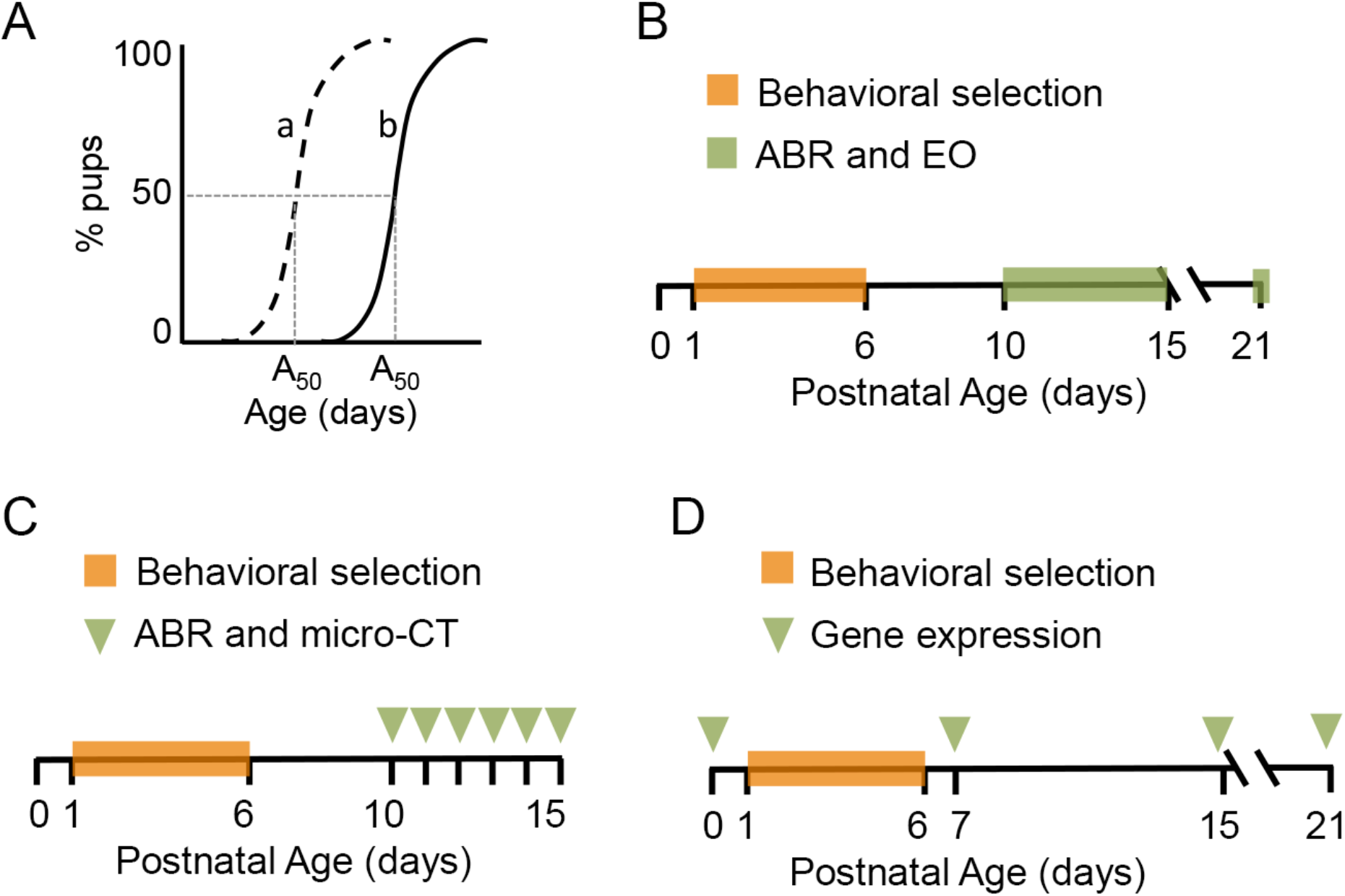
Experimental approach. **A,** The percent of pups with an auditory brainstem response (ABR) or eye opening (EO) in a litter increases as a function of age. Fitting such data to equation 1 determines an A_50_ value, the age at which 50% of pups in a litter show an ABR or EO. In this study we test the hypothesis that variation in maternal licking and grooming (LG) is associated with pup’s early (dashed line a) and late (continuous line b) sensory development profiles. **B,** Four maternal selection experiments were performed during spring and summer seasons (2 replicates per season, with cohorts of 40 dams per experiment). After dams gave birth (P0) maternal behavior was scored daily from P1 to P6 (n=135 dams). ABRs were tracked daily in all pups from seventeen selected litters between P10 and P15, and at P21. EO was tracked daily between P10 and P21 (n=199 pups). **C,** After behavioral scoring and selection between P1 and P6, correlative ABR and micro-CT X-ray tomography measurements were obtained from two low-LG litters (n=14 pups) and from two high-LG litters (n=15 pups) between P10 and P15 (indicated by arrowheads). **D,** Brain samples were obtained at different ages for gene expression analysis, prior and after behavioral scoring from three low-LG litters and four high-LG litters at P7, P15, and at P21 (indicated by arrowheads). Three litters were used to collect pups at P0, which was defined as a baseline for expressing fold changes in gene expression at other ages (n=3 pups per age per LG condition).

## METHODS

### Experimental Design and Statistical Analyses

Four maternal selection experiments were performed during the spring and fall seasons of two consecutive years (two experiments per year). For every selection experiment a cohort of 20 male and 40 female Wistar rats at postnatal day 65 was obtained from a commercial supplier (Charles River). From 160 females used in the study, a total of 137 had successful pregnancies (range of 32 to 36 dams per cohort). A total of 199 pups from 17 selected litters were used for developmental tracking. These included 81 pups from 7 low-LG litters (36 females and 45 males) and 118 pups from 10 high-LG litters (68 females and 50 males). Low-LG litters had an average of 12 ± 3 pups (mean ± SD; range 6 to 16 pups; n=7 litters). High-LG litters had an average of 12 ± 3 pups (mean ± SD; range 6 to 16 pups; n=10 litters). In addition, a total of 56 pups from four litters were used for correlative ABR and micro-CT measurements. These included 27 pups from two low-LG litters (11 females and 14 males) and 29 pups from two high-LG litters (11 females and 18 males) that were obtained from the first and fourth cohorts (one low-LG litter and one high-LG litter from each cohort). The expression of 30 genes implicated in the development and physiology of neuronal, glial and vascular cells in brainstem and cortical brain regions was examined with qRT-PCR in a total of 21 pups from either sex obtained from spring cohorts. Target genes were analyzed in three broad groups: a) neurotrophins, transcription factors and signaling effectors; b) oligodendrocyte development, hypoxia-sensitive pathway and mTor/Wnt signaling; and c) membrane transport. We compared 4 developmental stages comprising birth (P0; n=3 pups, each from different litters); the end of the first postnatal week at P7 (n=3 low-LG pups from two litters, and 3 high-LG pups from one litter); the end of the second postnatal week at P15 (n=3 low-LG pups from two litters, and 3 high-LG pups from one litter); and the weaning age at P21 (n=3 low-LG pups, each from different litters, and 3 high-LG pups from two litters).

Unless indicated, data represent mean ± SD. Statistical analyses were done with Prism 6 software (GraphPad). When appropriate, data sets were tested for normality using the D’Agostino and Pearson omnibus K2 test. Means in **Figure 2** were compared with an ordinary one-way ANOVA and the Holm-Sidak’s multiple comparisons test (**Figure 2A**), or the Tukey’s multiple comparisons test (**Figure 2D**). Medians in **Figure 3** were compared with the ANOVA Kruskal-Wallis test and the Dunn’s multiple comparisons test (**Figure 3E and F**). Gene expression data was analyzed by ANOVA Kruskal-Wallis test and the Dunn’s multiple comparisons test. Software built in algorithms for adjusting P values for multiple comparisons were used. Alpha = 0.05 was used to denote significance when testing for statistical differences between means or medians.

**Figure 2.**
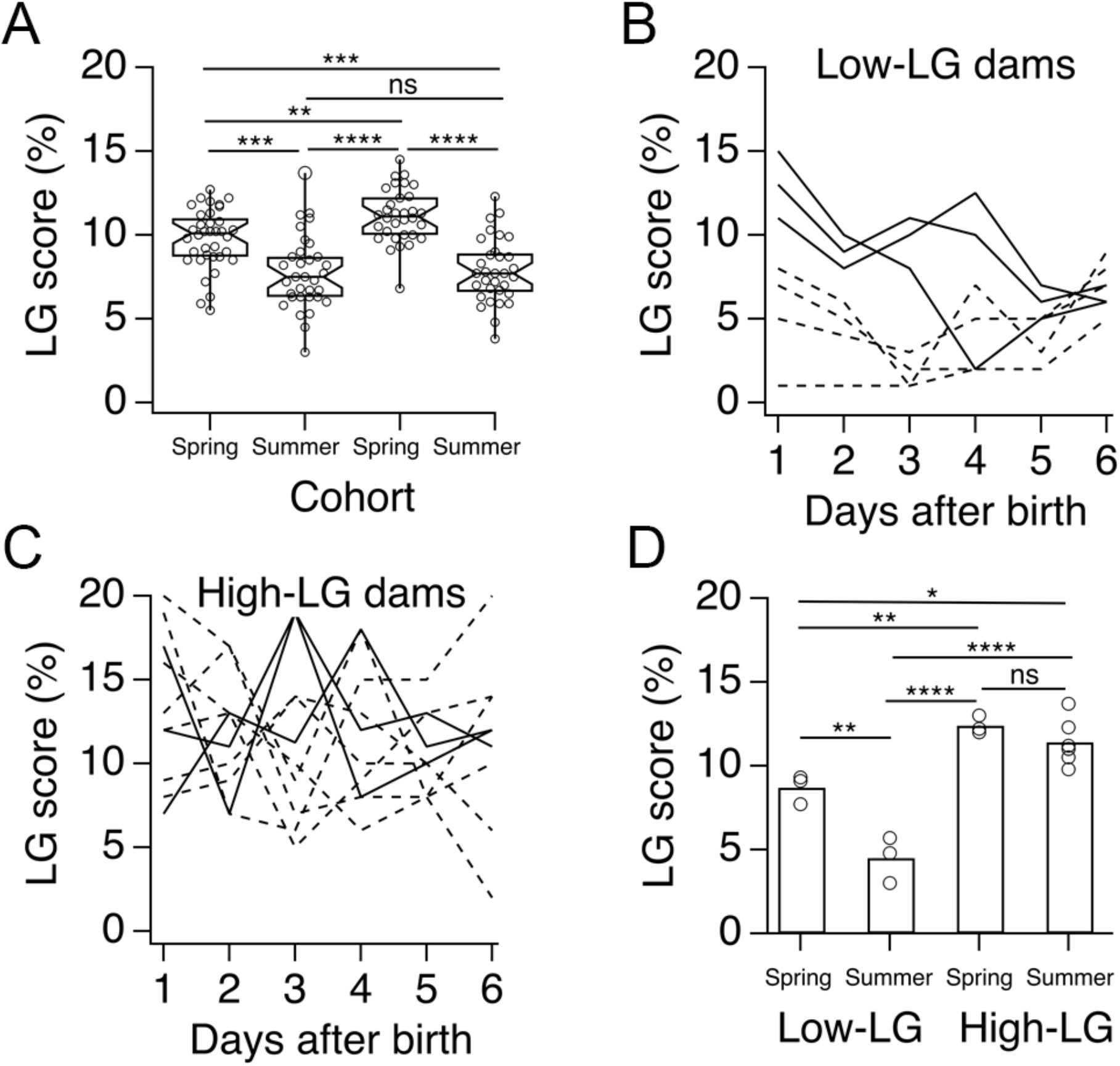
Variation in licking and grooming (LG) across different dam cohorts. **A,** Box plots of six-day average LG scores from cohorts analyzed in the spring and summer of two consecutive years. Each circle represents one dam. Asterisks indicate P values obtained by Holm’s Sidak’s multiple comparisons test, **=0.0021, ***=0.0001 and 0.0005, ****<0.0001. **B,** Daily LG scores for seven low-LG dams selected from spring and summer cohorts (continuous lines and dashed lines, respectively). **C,** Daily LG scores for ten high-LG dams selected from spring and summer cohorts (continuous lines and dashed lines, respectively). **D,** Six-day average LG scores for low LG and high-LG dams selected from spring and summer cohorts. Each circle represents one dam. Asterisks indicate P values obtained by Tukey’s multiple comparisons test, *=0.0249, **=0.0035 and 0.0088, ****<0.0001. n.s.=not significant.

### Animal housing and breeding

Experiments were performed in accordance with the Institutional Animal Care and Use Committee of the City College of New York. Rats were kept in a controlled environment at 22°C with an alternating 12 h light and dark cycle (lights were on at 7:00 hrs and off at 19:00 hrs). Water and food were available *ad libitum*. Male and female Wistar rats at postnatal day 65 were obtained from a commercial supplier (Charles River). Upon arrival, same sex rat pairs were housed in Plexiglas cages and acclimated to the animal care facility for one week. After acclimation, simple randomization with shuffled cage numbers was used to assign single males to a cage with a female pair. Breeding trios were housed together for five days. At the completion of the breeding period males were removed from the study and female pairs were housed together for 14 more days. Wistar rats have a gestation period of 22 days. Hence, 19 days after mating females were housed individually in Plexiglas cages that were supplied with paper towels as nesting material. Cages were checked for births everyday at 9:00 hrs, 12:00 hrs, and 17:00 hrs. On the day of birth (P0), pups were weighted and dams and their litters were placed in clean Plexiglas cages. Females that were not pregnant were removed from the study. Cages were undisturbed during behavioral scoring between P1 and P6, and routine twice per week cage cleaning resumed after behavioral scoring was finished. At P8, dams and their litters were transported to a satellite room for acclimation before testing.

### Maternal behavior scoring and selection criteria

Methods for scoring maternal behavior and litter selection were adapted from a previous study (Champagne et al., 2003). In brief, five 1-hour maternal behavior observation sessions were performed daily at 6:00 hrs, 9:00 hrs, 13:00 hrs, 19:00 hrs, and 21:00 hrs between pup ages P1 to P6. Every 1-hour observation consisted of 3 minute-long bins where the following behaviors were scored if observed: no contact with pups, contact with pups, dam is drinking, dam is eating, dam is self-grooming, dam is nest building, dam is licking and grooming (LG) pups in the anogenital or body region, and various levels of arched-back nursing described previously (Champagne et al., 2003). Unless indicated, LG scores in this study represent the frequency of LG in 100 observations per day (60 observations in the light cycle and 40 observations in the dark cycle) expressed as percent LG per day, or as average percent LG obtained from six-day scores. For every cohort, LG histograms were generated and individual dams were selected if their six-day average LG score was 1 SD above (high-LG) or 1 SD below (low-LG) the six-day average LG score of their cohort. Dams and litters that were not selected were removed from the study.

### Developmental tracking

Gender, body weight, onset of auditory brainstem responses (ABRs) and eye opening (EO) were tracked. Pup weight was recorded at P0, daily between P10 and P15, and at P21. Pup gender was determined between P10 and P15 using as joint criteria the anogenital distance and the presence or absence of multiple nipples to distinguish between males and females. EO was determined between P10 and P21, and was scored if at least one eyelid was open.

### Auditory brainstem response (ABR) tracking

All ABR measurements were done blind to LG group. ABRs were obtained daily between P10 and P15, and at P21. Anesthesia was induced inside a Plexiglas chamber with 3-5% isoflurane and maintained through a nose cone with 1.5% isoflurane dissolved in medical grade oxygen (gas flow set at 1 L min^−1^). ABRs were performed inside a double wall sound attenuated room (IAC). Anesthetized pups were placed onto a heating pad set at 37°C to keep them warm throughout the procedure. Subdermal electrodes were placed behind the right ear (reference electrode), at the vertex (active electrode), and at the left shoulder (ground electrode). A calibrated electrostatic Kanetec MB-FX free field speaker was used to deliver click sounds at 40 Hz with intensities ranging from 102 to 2 dB sound pressure level (SPL) in 5 dB decrements. Clicks were synthesized with TDT system 3 hardware (Tucker-Davis Technologies), and presented at 20 kHz with alternating polarity to minimize the presence of stimulus artifacts. Speaker calibration was done with a type 7012 ½ inch ACO Pacific microphone (reference 20 μPa). ABR waveforms were recorded with a Medusa preamplifier at 24.4 kHz and saved to hard disk for offline analysis (Tucker-Davis Technologies). ABRs in this study are average waveforms of 300 traces with 10 ms duration. ABR measurements per litter were completed in 30-40 minutes, including the time it took for pups to recover from anesthesia. After recovery, all pups were placed back into their home cages until the next day of testing.

### Combined ABR and micro-CT X-ray tomography (micro-CT) experiments

On the first day of experiments at P10, pups within a litter were labeled with permanent ink, sexed and processed for ABRs in pairs. After ABRs were measured, anesthetized pups were decapitated and their heads were processed fresh for micro-CT imaging. Micro-CT images were acquired and processed as described previously (Adise et al., 2014). X-Ray projections were generated around the samples with 0.4° rotation steps at a resolution of 11.5 **μ**m per pixel using a 1172 Bruker SkyScan (Bruker). Scans were loaded into MIMICS (v14.0, Materialise) for segmentation and 3D reconstruction. With the exception of the postalignment compensation, all reconstruction parameters were applied identically to all scans. Micro-CT imaging was performed blind to LG group. This procedure was repeated between P10 and P15 until all pups within a litter were used. For the low-LG and the high-LG litters obtained from the last selection experiment, all pups were screened for ABRs between P10 and P15 and pairs were removed daily for micro-CT imaging. This procedure allowed us to track ABRs between P12 and P13, when major changes in air volume of middle ear cavity and physiological responses were observed.

### Gene Expression

qRT-PCR was performed with QuantStudio 7 Flex Real-time qPCR system (Thermofisher), using protocols available at the Advanced Science and Research Center Epigenetic Core Facility. Briefly, primer pairs were obtained from a commercial vendor (Sigma-Aldrich) and primer specificity was tested with adult rat whole-brain cDNA. **Table 1** shows the list of primers used in this study in the same order as they appear in **Figures 6, 7 and 8**. Total RNA was isolated from a bank of frozen brains kept at −80 °C using a RNA isolation kit according to the manufacturer’s instructions (Qiagen). Frozen brain samples were thawed and dissected from 5 different regions: cochlear nucleus, pons (ventral brainstem containing the acoustic stria), inferior colliculus, temporal cortex (here referred as auditory cortex), and occipital cortex (here referred as visual cortex). Reverse transcription and specific target amplification were completed using qScript cDNA Supermix (Quanta) according to manufacturer’s protocol. A primer mixture containing both forward and reverse primers was mixed with cDNA from different brain regions and loaded onto 384 well plates. The QuantStudio analysis software was used for data analysis and visualization. Threshold was determined automatically and Ct values were calculated using QuantStudio analysis software. The housekeeping gene *Actb* (coding for β-actin) was measured for all ages and LG conditions tested using primers described in **Table 1**.

**TABLE 1.**
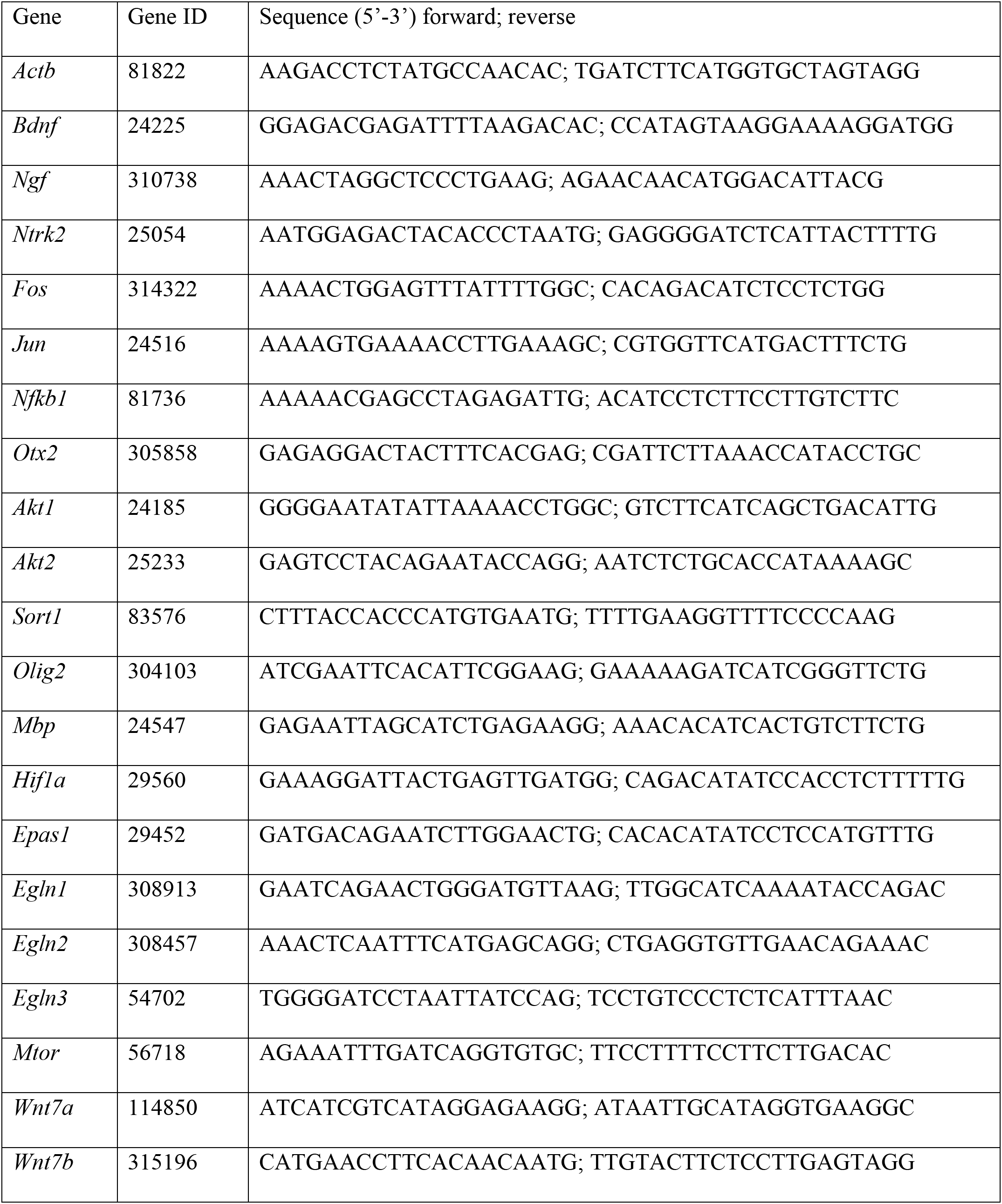
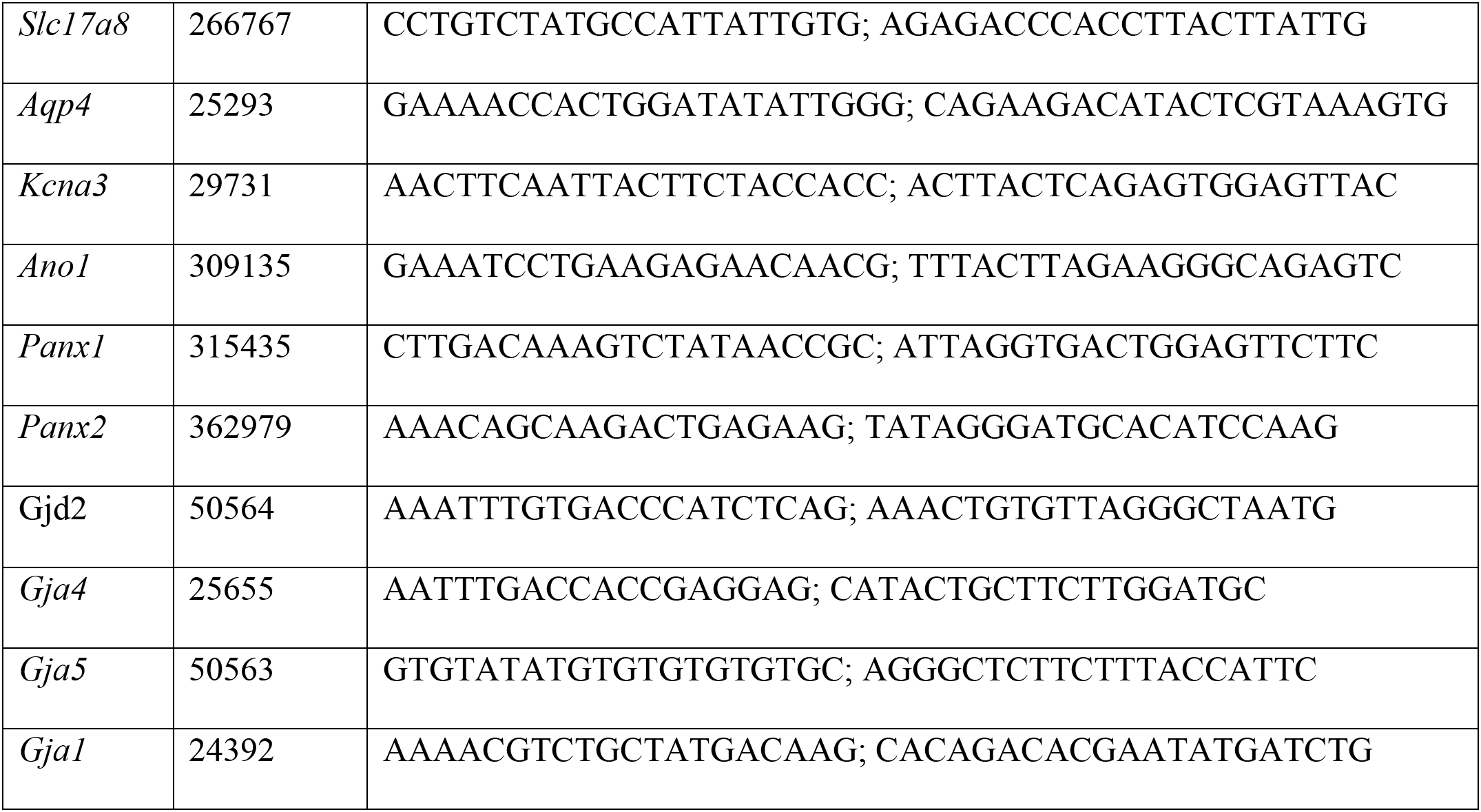
List of primer pairs used for analysis of gene expression with qRT-PCR.

### Data Analysis

ABR recordings were saved as text files and analyzed using NeuroMatic in Igor Pro software (WaveMetrics; Rothman and Silver, 2018). ABR thresholds were determined using an amplitude criterion to detect responses that were larger than four times the standard deviation (SD) of the baseline (Bogaerts et al., 2009). In general, waveforms with amplitudes larger than 1 microvolt were considered auditory responses. Wave I was defined as a positive transient voltage change with a peak latency of 1.8 ms to 2.2 ms. Short latency potentials (SLPs) were defined as positive transient voltage changes with a peak latency ~ 1 ms.

Developmental curves of percent pups with a wave I response, EO, or air volume at different ages were fit to **Equation 1**:

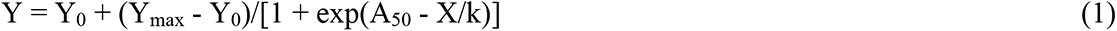

Where Y_0_ is the minimum observed Y (i.e., the percent pups with ABR, EO, or air volume), Ymax is the maximum observed Y, A_50_ is the age at which Y is half maximum, X is age (in days), and k is the rate coefficient.

## RESULTS

### Variation in maternal LG

We used four Wistar rat cohorts from the spring and summer seasons of two consecutive years to select low-LG and high-LG dams, and we tracked the sensory development of their pup’s between postnatal ages P10 and P21 (**Figure 1B**). **Figure 2A** shows box plots of maternal LG scores from the four cohorts used in this study. The six-day average LG score for each cohort was (mean ± SD): 9.8 ± 1.8 (n=36 dams, spring year 1); 7.7 ± 2.2 (n=33 dams, summer year 1); 11.2 ± 1.6 (n=36 dams, spring year 2); and 7.9 ± 1.9 (n=32 dams, summer year 2). Statistical analysis showed significant differences between mean LG scores (Ordinary one-way ANOVA, F=25.91, P<0.0001), in particular between spring cohorts, and between spring and summer cohorts. Mean summer cohort LG scores were not significantly different from each other (determined by Tukey’s multiple comparisons test; see **Figure 2** legend for P values). The large variability in LG scores across cohorts prompted us to examine the daily LG scores of the seven low-LG dams and the ten high-LG dams that were selected. Low-LG dams selected from spring cohorts showed daily LG profiles that started high and decreased during the six-day observation period (continuous lines in **Figure 2B**), while low-LG dams selected from summer cohorts showed relatively lower LG scores throughout the six-day observation period (dashed lines in **Figure 2B**). High-LG dams had daily LG scores that were very variable but stayed relatively high throughout the six-day observation period, regardless of whether they were obtained from the spring or summer cohorts (**Figure 2C**). Statistical analysis showed significant differences between the six-day average LG scores of selected dams (**Figure 2D**; ordinary one-way ANOVA, F=31.93, P<0.0001). We found that the average LG score of low-LG dams was higher in spring cohorts compared to summer cohorts (Tukey’s multiple comparisons test, P=0.0035). In contrast, we did not find statistically significant differences between the six-day average LG scores of high-LG dams from spring and summer cohorts (Tukey’s multiple comparisons test, P=0.5672). Overall, these results show that despite the variable LG scores between cohorts, the LG scores of selected low-LG and high-LG groups were significantly different from each other.

### Variation in auditory brainstem response (ABR) onset and eye opening (EO) in pups reared by low-LG and high-LG dams

ABRs and EO were tracked in a total of 81 pups from seven low-LG litters and in 118 pups from ten high-LG litters. For each litter the percent of pups with an ABR wave I, or the percent of pups with EO were plotted at different ages, and fits to **Equation 1** were obtained (**Figure 3A, B, C and D**; continuous lines represent fits to data from spring litters, dashed lines represent fits to data from summer litters). To examine the variation in ABR onset and EO within and across LG groups, the distributions of A_50_ values were compared (**Figure 3E**). This qualitative analysis showed skewed ABR A_50_ distributions for low-LG and high-LG litters, indicating that the majority of pups examined had early ABR onset between P11.5-P12. However, in some litters pups had ABR onset as late as P13-P13.5. EO A_50_ distributions for low-LG and high-LG litters were also skewed and covered a range of two days between P13 and P15. Statistical analysis showed significant differences between A_50_ medians (Kruskal-Wallis test, P value <0.0001). A more detailed examination showed that the ABR A_50_ medians between low-LG and high-LG litters were not significantly different from each other (Dunn’s multiple comparisons test, see **Figure 3** legend for P values). A similar result was obtained for EO A_50_ medians (Dunn’s multiple comparisons test, see **Figure 3** legend for P values). However, the ABR A_50_ medians were significantly different from the EO A_50_ medians, within and across LG groups (indicated by asterisks in **Figure 3E**; Dunn’s multiple comparisons test, see **Figure 3** legend for P values). To obtain information on the synchrony of development within litters, ABR and EO rate coefficient k distributions were compared (see **Equation 1** and **Figure 3F**). The data shows evidence of predominately short values of rate coefficient k for low-LG and high-LG litters. Neither ABR rate coefficient k medians nor EO rate coefficient k medians showed significant differences within and across LG groups (Kruskal-Wallis test, P value=0.1171). In sum, experiments described in **Figures 1B, 2,** and **3** show that despite significant differences in LG scores between selected dams, ABR onset, timing of EO and synchrony of development do not differ between low-LG and high-LG litters. Instead, there is a range of litter-specific early or late times for ABR onset that happens during a two-day period. Similarly, a two-day range for early and late EO takes place sequentially after ABR onset.

**Figure 3.**
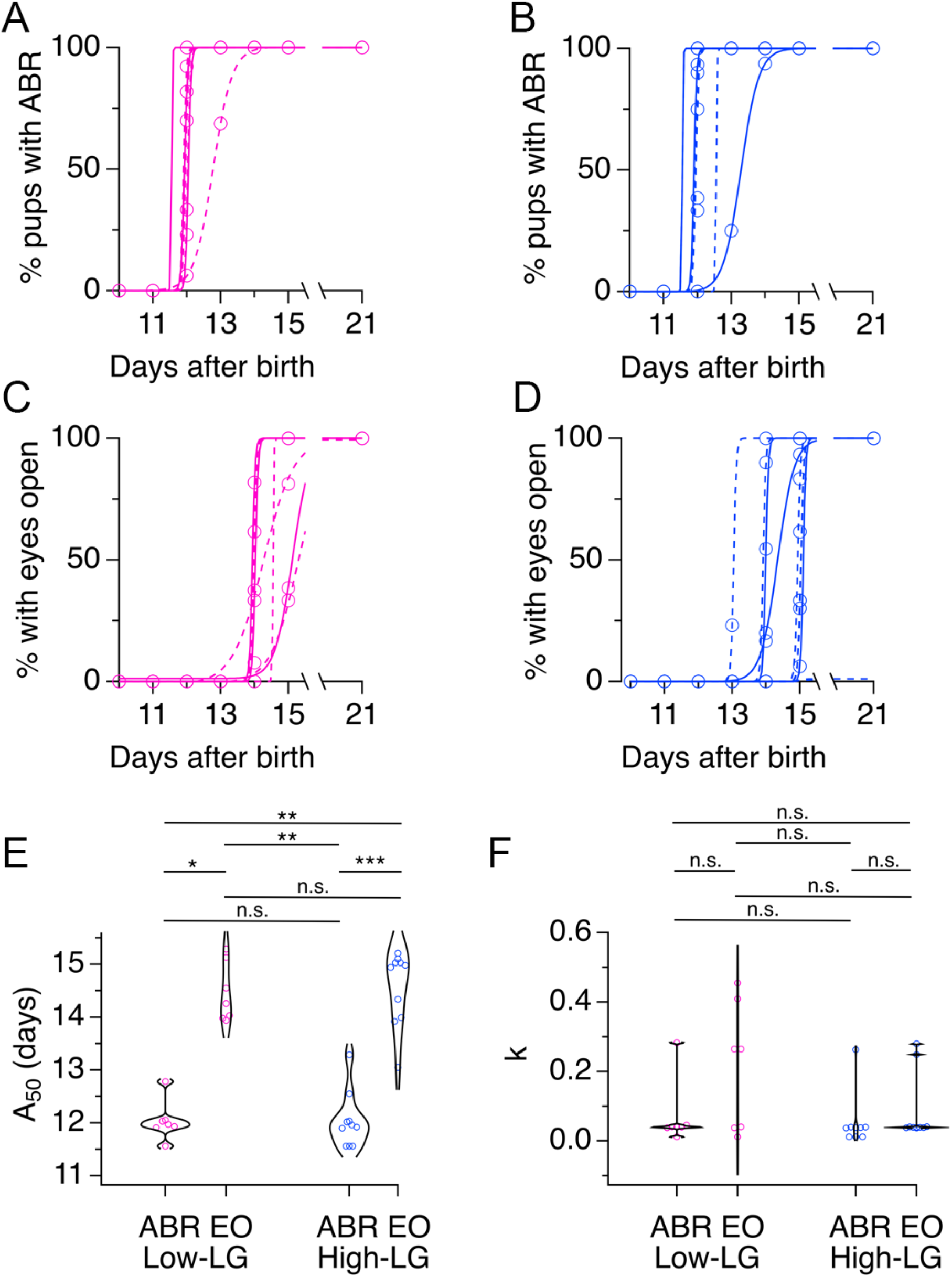
Timing of auditory brainstem response (ABR) onset and eye opening (EO) in the offspring of selected dams. **A,** Plots of percent pups with ABR wave I at different ages from seven low-licking and grooming (low-LG) litters were fit to equation 1. **B,** Plots of percent pups with ABR wave I at different ages from ten high-licking and grooming (high-LG) litters were fit to equation 1. **C,** Plots of percent pups with EO at different ages from seven low-LG litters were fit to equation 1. D, Plots of percent pups with EO at different ages from ten high-LG litters were fit to equation 1. **E,** Violin plots of A_50_ values obtained from fits of equation 1 to developmental data of percent pups with ABR or EO. **F,** Violin plots of rate coefficient k values obtained from fits of equation 1 to developmental data of percent pups with ABR or EO. Asterisks indicate significant differences between medians: * indicates P value = 0.0172; ** indicate P values = 0.0055 and 0.0027; *** indicates P value =0.0005; Dunn’s multiple comparisons test; n.s.=not significant. Continuous lines represent fits to spring litters; dashed lines represent fits to summer litters.

### Maternal LG is not correlated with ABR onset or EO

We used scatter plots of ABR A_50_ or EO A_50_ values against LG scores to determine the correlation coefficients between these two variables. We did not find evidence of a correlation between ABR A_50_ values and LG scores (non parametric Spearman correlation, R=−0.1063, two-tailed P=0.6823), or between EO A_50_ values and LG scores (non parametric Spearman correlation R−-0.05184, two-tailed P=0.8434). We also checked for systematic differences in the growth of pups reared by low-LG and high-LG dams. We compared average pup body weight between females and males within a litter and across LG groups on the day of ABR onset (defined by the A_50_ parameter) or using the slope of the growth curve between P10 and P15 (determined by linear regression of body weight data). This analysis showed that pup’s body weight did not correlate with maternal LG scores. Similar results were obtained when we tested for developmental differences between male and female pups (data not shown). Overall, the data discussed in this section does not support a correlation between maternal LG scores and developmentally tracked features of male and female pups.

### Differences in the delay between ABR onset and EO in litters with early or late ABR onset

Next, we examined ABR onset and EO data in scatter plots of EO A_50_ values plotted against ABR A_50_ values (**Figure 4A**). This analysis showed that in individual litters, ABR onset always happened before EO, and notably, it confirmed that for both, ABR and EO, there was a range of early and late onset times that happened within a two-day window: while ABR onset was observed between P11.5 and P13.5, EO was observed between P13 and P15. Note that there was never a litter in which the age of EO coincided with the age of ABR onset. Instead, litters with the earliest ABR onset had more variable EO times than litters with late ABR onset. In litters with ABR onset around P11.5 and P12, we observed delays to EO from 1.5-days to 3.5-days. In contrast, in litters with late ABR onset at ~P13, there was a ~1.5-2 day delay to EO. This pattern was observed in low-LG and high-LG litters alike (**Figure 4A**). A similar comparison between EO and ABR rate coefficient k showed that in most litters ABR rate coefficient values were <0.1, implying developmental synchrony within litters. In contrast, EO rate coefficients were more variable, implying developmental synchrony and asynchrony, respectively across different litters (**Figure 4B**).

**Figure 4.**
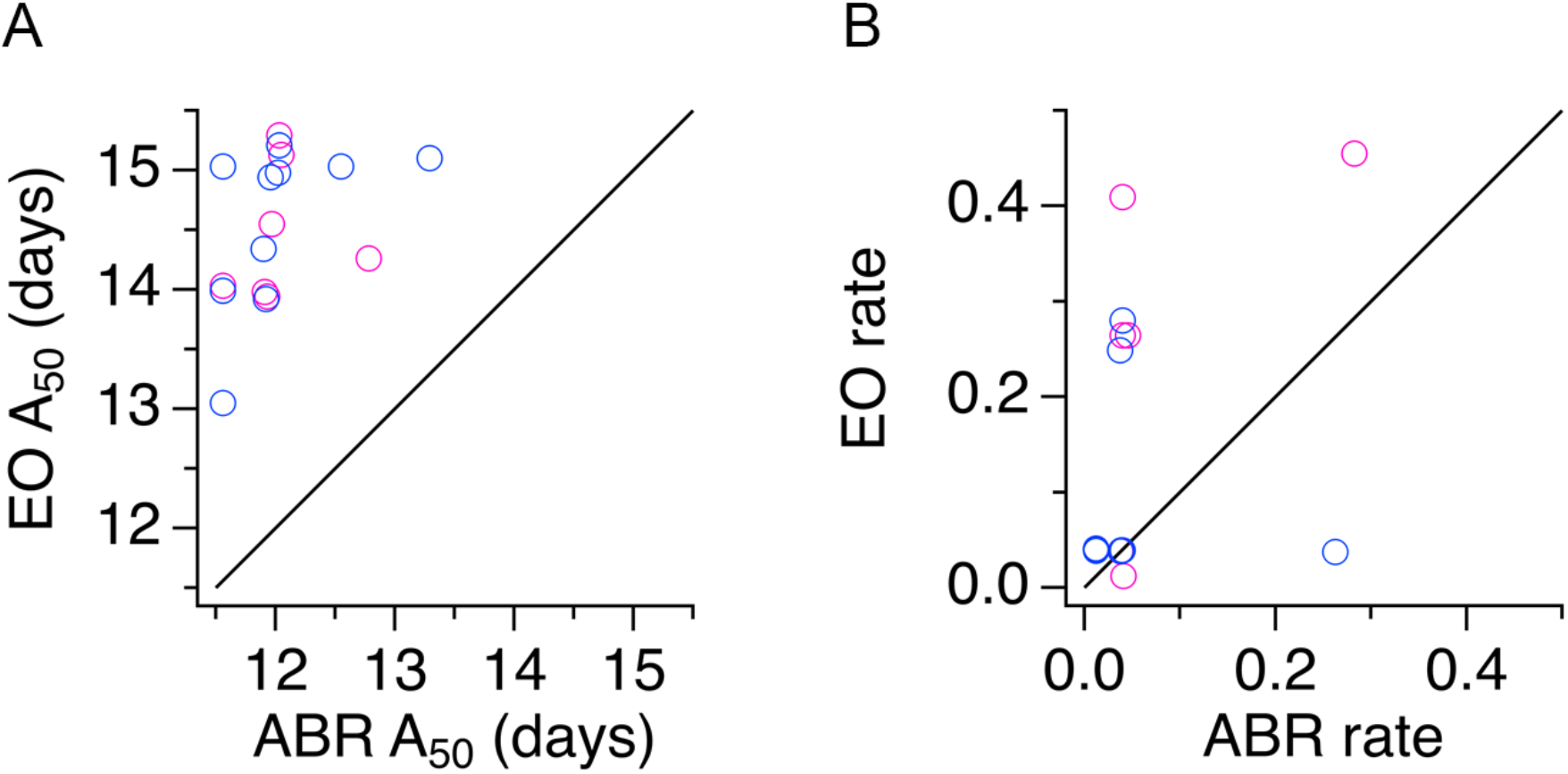
Delay between auditory brainstem response (ABR) onset and eye opening (EO). **A,** Scatter plot of A_50_ values show the relationship between EO and ABR onset in low-licking and grooming (low-LG) and high-licking and grooming (high-LG) litters. **B,** Scatter plot of rate coefficient k values show the relationship between EO and ABR development in selected low-LG and high-LG litters. Magenta symbols represent low-LG data; blue symbols represent high-LG data. Black line in A and B represents the identity line.

### Relationship between the development of the middle ear and ABR thresholds in the progeny of low-LG and high-LG dams

To obtain information about developmental structural changes in the auditory periphery of pups from low-LG and high-LG litters, four litters were used to perform correlative ABR and micro CT X-ray tomography (micro-CT) experiments (**Figure 1C**; n=51 pups). Detailed examination of the 3D renderings generated from micro-CT data confirmed our previous finding that formation of the middle ear cavity precedes formation of the ear canal (**Figure 5A**; Adise et al., 2014). However, in contrast to previous studies, precursor zones or small air pockets were not observed. Instead, there were marked differences in the air volume of pups from different litters, particularly between P12 and P13 (**Figure 5B**). Fitting **Equation 1** to data in **Figure 5B** gave A_50_ values that ranged from 11.8 days to 13.2 days (12.4 ± 0.3 days, n = 4 litters), and rate coefficient k values that ranged from 0.28 to 0.47 (0.40 ± 0.04, n = 4 litters). From the 51 pups used in micro-CT imaging experiments, we confirmed an ABR wave I in 25 pups between P12 and P15, while 17 pups between ages P10 and P11, and 9 pups between ages P12 and P13 did not show any evidence of ABR wave I (**Figure 5C**). Since the micro-CT imaging did not detect air in any of the 17 non-responsive (NR) pups examined between P10 and P11, we can infer that formation of an air-filled middle ear cavity is necessary for transmission of airborne pressure waves to the inner ear. Linear fitting of wave I threshold data versus air volume in the range between 25 mm^3^ to 60 mm^3^ gave a slope of −2.5 dB/mm^3^, showing that auditory thresholds are inversely proportional to air volume in the auditory periphery. However, the structural data also suggests that the presence of air in the middle ear may not be sufficient for proper sound transmission, since there were 7 animals with a measurable air volume in the middle ear at P12 and P13 that did not show an ABR wave I (labeled non responsive, NR, in the boxed area of **Figure 5C**).

**Figure 5.**
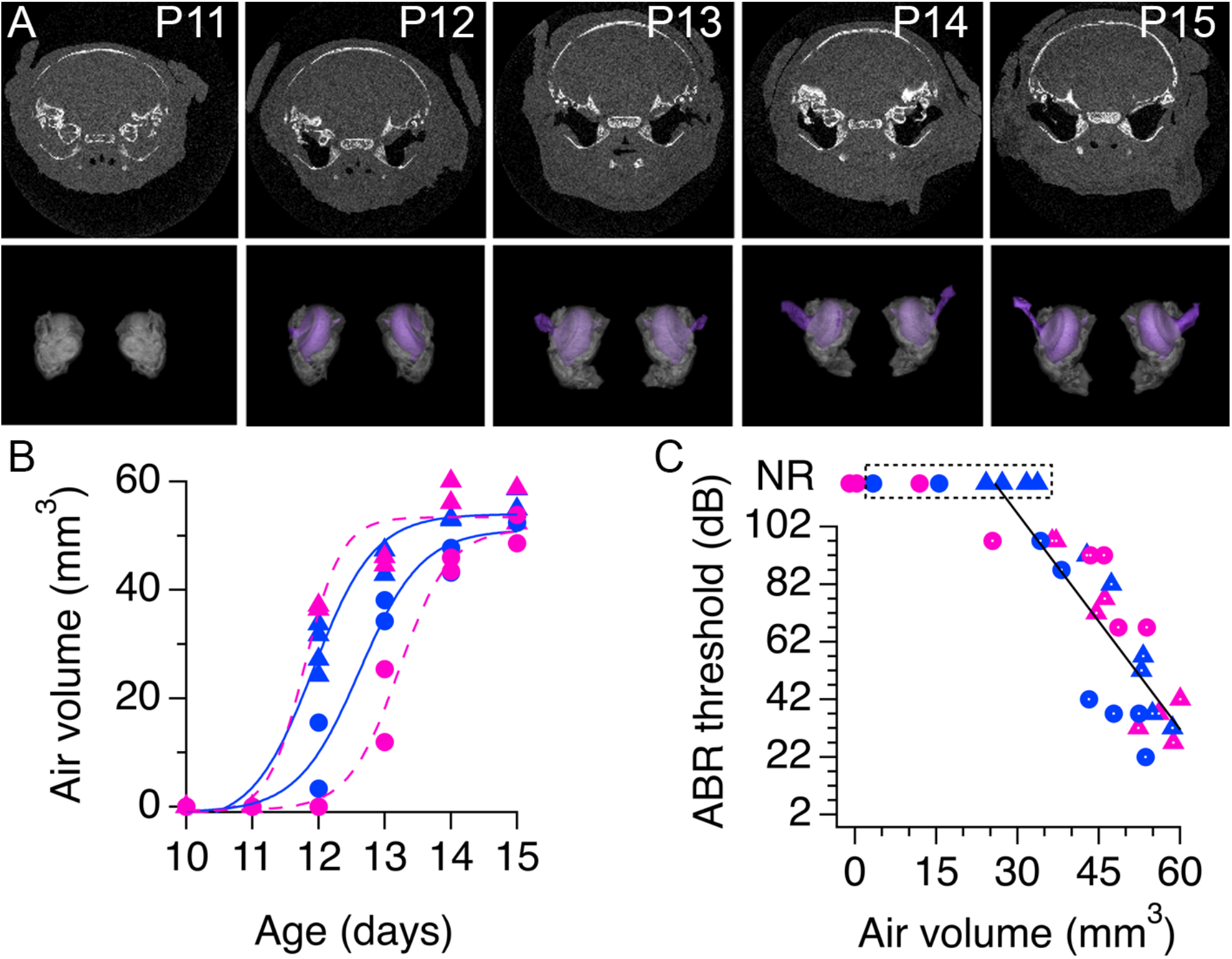
Relationship between development of the middle cavity and wave 1 auditory brainstem (ABR) thresholds. **A,** Top, developmental series of micro-CT X ray scans of pups from a low-licking and grooming (low-LG) litter. White indicates bone, grey is soft tissue and black is air. Bottom, 3D rendering of segmented bone (gray) and air (purple) contrast obtained from tomographic data. **B,** Air volume measured at different ages in pups from two low-LG (magenta) and two high-licking and grooming (high-LG) litters (blue). Note that every symbol represents one pup and that similar symbols represent pups from the same litter. Continuous and dotted color lines are fits of equation 1 to the data. **C,** Relationship between ABR wave I thresholds and air volume in the middle ear cavity. Magenta circles and triangles represent pups from low-LG litters (n = 25 pups); blue circles and triangles represent pups from high LG litters (n = 26 pups). NR = non-responsive pups, defined by the absence of ABR wave I. Black line represents the fit to a linear function between wave I thresholds and air volume with slope 2.5 dB/mm^3^. Symbols with white dots indicate data points included in the fit.

To examine the possibility that a minimal air volume at the auditory periphery is necessary for the onset of ABRs, the ABR waveforms from all pups at P12 (n=10) and all pups at P13 (n=8) used in the combined ABR and micro-CT experiments were re-examined. To our surprise, seven P12 pups and six P13 pups with air volumes larger than 12 mm^3^ had responses of comparable amplitude to wave I, but with a shorter latency. We refer to these events as short latency potentials (SLPs; **Figure 6**). **Figure 6D** shows exemplar ABR traces with SLPs at different click intensities in a P12 pup whose structural information is shown in **Figure 6C**. Note that in this example wave I was not present, determined by the absence of a positive potential with a latency ~2 ms. **Figure 6F** shows exemplar recordings from another P12 pup that had SLPs followed by wave I at different click intensities and whose structural data is shown in **Figure 6E**. Note that in this example a wave I was identified after the SLP at click intensities of 102 dB and 97 dB but not at lower intensities, demonstrating that the threshold for the SLP was lower than the threshold for wave I. Lastly, exemplar ABR traces are shown for a P12 pup with an air volume of zero in the middle ear. In this case SLPs and wave I were absent in response to the same click intensities probed for the other pups (**Figure 6A and B**). Based on these observations, we re-examined all the ABR recordings from the 4 litters used in combined ABR and micro-CT experiments between P11 and P15 to corroborate the presence or absence of SLPs at different ages. We found evidence of SLPs at P12 (n = 7 pups) and P13 (n = 6 pups), but we did not find any evidence of SLPs at P11 (n= 8 pups), P14 (n = 8 pups) and P15 (n = 8 pups). **Figure 6G** plots the SLP thresholds as a function of air volume for all ten P12 and eight P13 pups used in the combined ABR and micro-CT experiments (**Figure 6G**, solid symbols). Note that there were four NR P12 pups whose air volumes were < 15 mm^3^ and did not have SLPs or ABR wave I, and one P13 NR pup with an air volume of 34 mm^3^ that did not have a SLP but had an ABR wave I (P12 NR pups are enclosed together in a dashed box, and the P13 NR pup with wave I but without SLP is enclosed in a dashed box marked with an arrow in **Figure 6G**). Linear fitting of SLP threshold data versus air volume in the range between 15 mm^3^ to 50 mm^3^ gave a slope of − 0.3 dB/mm^3^. Altogether, these data support the view that a minimal air volume at the auditory periphery is necessary for airborne conduction of click sounds from the external ear to the inner ear. Next, we examined the relationship between SLPs and wave I responses.

**Figure 6.**
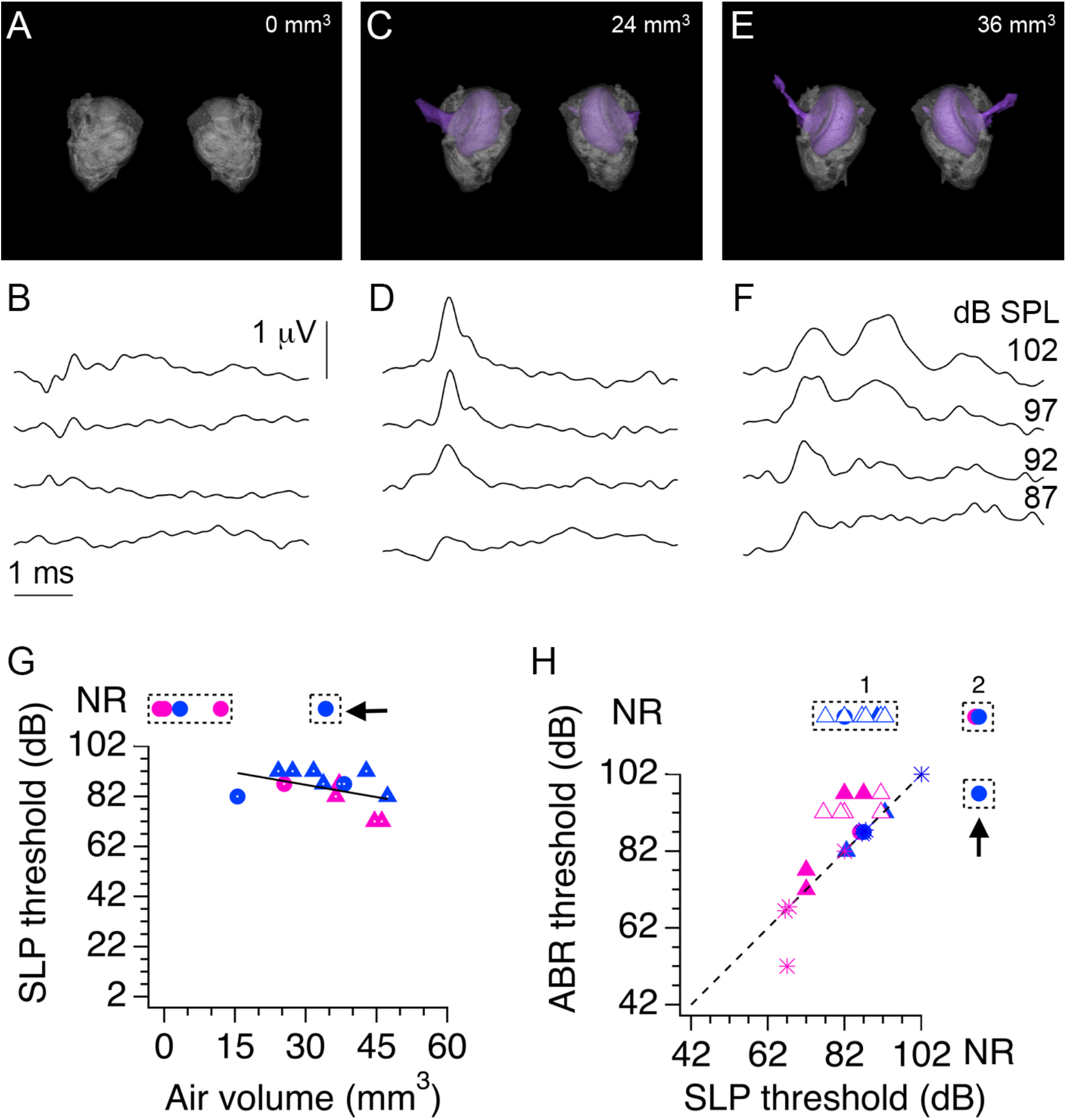
Identification of short latency potentials (SLPs) in combined auditory brainstem response (ABR) and micro-CT X ray (micro-CT) experiments. **A, C, E,** Representative 3D rendering of three P12 pups with different volumes of air in the middle ear and external canal. **B, D, F,** Corresponding ABR waveforms show the presence of a SLP in pups with air volumes of 24 mm^3^ and 36 mm^3^, but not in the pup without middle ear and external ear cavities. ABR traces in B, D and F correspond to click intensities shown in panel F. **G,** SLP thresholds as a function of air volume. Black line represents the fit to a linear function between SLP thresholds and air volume with slope 0.3 dB/mm^3^. Symbols with white dots indicate data points included in the fit. **H,** Comparison of SLP thresholds and wave 1 thresholds for pups measured at P12 and P13. Filled symbols represent pups used in combined ABR and micro-CT experiments. Open symbols are littermates used solely in ABR experiments at P12. Asterisks indicate pups tracked from P12 to P13. Black dashed line in H is the identity line. Arrows in G and H indicate a pup that showed ABR wave I but not a SLP.

### SLPs show hallmarks of sensory responses from the inner ear

We hypothesized that SLPs may represent electrical responses in hair cells of the inner ear. Alternatively, SLPs could represent evoked potentials from a different sensory modality, such as somatosensory fibers activated by the pressure energy contained in click stimuli of high intensity. If SLPs were generated in hair cells of the inner ear, then we would expect that SLPs and wave I would show hallmarks of synaptic communication, including a defined delay between events. In addition, we would expect developmental changes in the thresholds and the delay between SLPs and wave I. We would not expect to see these hallmarks if SLPs were sensory responses independent from wave I. To test these predictions, we took advantage that six littermates from one low-LG litter and six littermates from another high-LG litter were not used for micro-CT scans at P12. We recorded ABRs and defined the co-occurrence of SLPs and wave I, and examined how the threshold for SLPs changed with respect to the threshold of wave I between P12 and P13. We found that all six pups from the high-LG litter had SLPs but did not have a wave I (open blue triangles in **Figure 6H**). Interesting to us, five out of six pups from the low-LG litter had SLPs followed by a wave I, and one pup had SLPs without any evidence of wave 1 (open magenta triangles in **Figure 6H**). Counting all the pups used in combined ABR and micro-CT experiments and the subset of littermates used in ABR tracking we found that in low-LG litters at P12 there were 2 pups that did not have a SLP nor a wave I (**Figure 6H** box 2); 1 pup had SLPs but not a wave I; 6 pups had SLPs with thresholds that were lower than their corresponding wave I thresholds; and 1 pup had SLPs with a threshold that was similar to its wave I threshold. In high-LG litters at P12 there was 1 pup without SLP and wave I; 10 pups had SLPs and no wave I (**Figure 6H** box 1); and 1 pup had SLPs with a threshold lower than its wave I threshold. Thus, based on this data, it seems that SLPs occur alone at P12, and when SLPs and wave I are observed together, SLPs have lower thresholds than wave I. In low-LG litters at P13, there were 3 pups that had SLPs with thresholds that were lower than wave I thresholds. In high-LG litters at P13, all 3 pups had SLPs with thresholds that were similar to their corresponding wave I thresholds. Thus, at P13, SLPs always co-occur with wave I and had higher or similar thresholds than wave I.

To examine the development of SLPs and wave I responses, we tracked the ABRs from P12 to P13 in the eight remaining littermates from the low-LG and high-LG litters (4 pups per litter). We found evidence of a decrease in wave I thresholds from P12 to P13 such that in 7 of 8 pups SLPs had thresholds that were similar to their corresponding wave I thresholds, and in one pup the wave I threshold was lower than its corresponding SLP threshold (asterisks in **Figure 6H**). This single observation raised the possibility that in this animal, SLPs were independent events of wave I events. To test the possibility that somatosensory fibers could be activated by the pressure wave energy of high intensity click stimuli, we injected the local anesthetic lidocaine around the skin pad surrounding the pinna area in three P13 littermates used for ABR experiments. This manipulation did not affect the occurrence of SLPs in these animals, suggesting that skin stimulation by high intensity clicks does not generate SLPs (data not shown). Altogether, these results indicate that SLPs antecede the developmental expression of wave I responses (open symbols in **Figure 6H**) and suggest that as pups mature, wave I thresholds decrease to match SLP thresholds.

Lastly, we obtained estimates of two physiological parameters of SLPs: the latency of events from stimulus onset, and the delay between the peaks of SLP’s and wave I events at P12 and P13. The latency from stimulus onset was evaluated from ABR traces at different intensities ranging between 77 dB and 102 dB. We did not find systematic changes in this parameter as a function of click intensity at P12 or P13, so we obtained an average SLP latency per pup from these combined measurements and obtained a grand average per LG group per age. At P12, the latency of SLPs was 1.08 ± 0.01 ms in low-LG pups (n=8 pups) and 1.09 ± 0.01 ms in high-LG pups (n=11 pups). At P13, the latency of SLPs was 1.09 ± 0.01 ms in low-LG pups (n=7 pups) and 1.07 ± 0.01 ms in high-LG pups (n=7 pups). The mean values of SLP latency for different LG groups and ages were not significantly different from each other (age P=0.6342; LG group P=0.6342; interaction between age and LG group P=0.1598, 2 way ANOVA). At P12, the delay between the peaks of the SLP and wave I was 1.01 ± 0.03 ms in low-LG pups (n=7 pups), and could not be determined in high-LG pups since they did not have a wave I at this age, or if they had a wave I it was not preceded by a SLP (boxed data point indicated with an arrow in **Figure 6G and H**). At P13, the delay between the peaks of the SLP and wave I was 1.03 ± 0.03 ms in low-LG pups (n=7 pups), and 1.02 ± 0.02 ms in high-LG pups (n=7 pups). Altogether, data in **Figures 5 and 6** show the relationship between development of the auditory periphery and the type of sensorineural response recorded in the progeny of low-LG and high-LG pups. SLPs predominated over wave I responses at P12, and gradually waned as wave I responses increased in amplitude at P13 and thereafter.

### Analysis of gene expression in the auditory brainstem, auditory cortex (ACX) and visual cortex (VCX) of pups reared by low-LG and high-LG dams

A gene expression screen using qRT-PCR was carried out with pup tissue from five brain regions at four ages. We screened the relative mRNA expression levels of 30 genes involved in neuronal, glial and vascular physiology and development in samples from the cochlear nucleus (CN), the pons, the inferior colliculus (IC), the primary auditory cortex (ACX), and the primary visual cortex (VCX) from neonate pups at P0, P7, P15, and P21 (Figure 1C; n=3 pups per age per LG group). Gene expression data is expressed as fold change with respect to P0 and summarized in **Figures 7, 8 and 9**.

**Figure 7.**
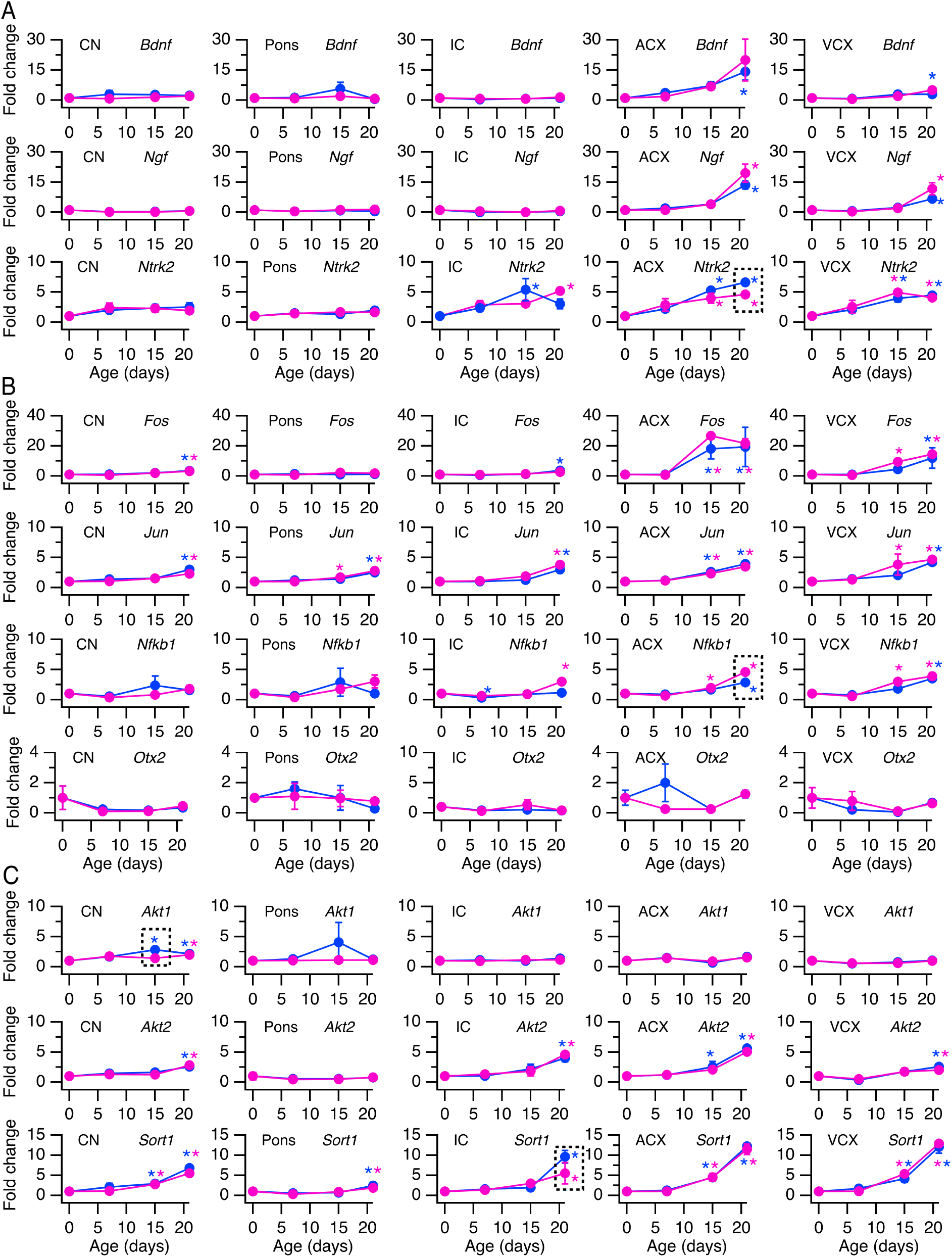
Temporal expression profiles for genes involved in neural development and plasticity. **A,** Relative level of mRNA expression of neurotrophin genes BDNF, NGF and the BDNF receptor TrkB. **B,** Relative level of mRNA expression for transcription factors c-Fos, c-Jun, NFκB and Otx2. **C,** Relative level of mRNA expression for signaling effectors Akt1, Akt2, and Sort 1. Data is plotted as fold change with respect to birth (P0). Magenta symbols represent data from low-licking and grooming (low-LG) samples. Blue symbols represent data from high-licking and grooming (high-LG) samples. Data represents mean ± sem (n=3 pups per age per LG group). Asterisks represent statistically significant differences with respect to P0. Boxed data represents statistically significant differences between low-LG and high-LG samples. Alpha=0.05.

### Analysis of genes involved in developmental plasticity

**Figure 7** shows results for 10 genes involved in developmental plasticity, including transcriptional regulation and signal transduction. **Figure 7A** shows expression profiles for neurotrophin genes *Bdnf* and *Ngf*, and *Ntrk2*, which codes for the Bdnf receptor TrkB. The relative levels of *Bdnf* mRNA in subcortical structures did not change between P0 and any other age examined (ordinary one-way ANOVA P=0.5135, P=0.5157, P=0.2458, for CN, pons, and IC, respectively). For cortical structures, there was a statistically significant increase in ACX and VCX of high-LG pups between P0 and P21 (ordinary one-way ANOVA P=0.0730, P=0.0175, for ACX and VCX, respectively; and multiple comparisons test P=0.0192 and P=0.0035, for ACX and VCX respectively). The relative levels of *Ngf* mRNA did not show statistically significant changes in subcortical structures between P0 and any other age examined (ordinary one-way ANOVA P=0.4500, P=0.4226, P=0.3039 for CN, pons and IC, respectively). However, *Ngf* mRNA levels showed a statistically significant increase in ACX and VCX of pups from both LG groups between P0 and P21 (ordinary one-way ANOVA P=0.0001 and P=0.0001, for ACX and VCX respectively; and multiple comparisons test P=0.0001, P=0.0007, for low-LG and high-LG samples in ACX respectively; P=0.0001, P=0.0007, for low-LG and high-LG samples in VCX, respectively). Similar to *Bdnf* and *Ngf* mRNAs, *Ntrk2* mRNA levels did not show changes in CN and pons between P0 and any other age examined (ordinary one-way ANOVA P=0.5272 and P=0.523 for CN and pons, respectively). However, there was a significant increase of *Ntrk2* mRNA levels in the IC of high-LG pups between P0 and P15, and a significant increase of *Ntrk2* mRNA levels in the IC of low-LG pups between P0 and P21 (ordinary one-way ANOVA P=0.0324; and multiple comparisons test P=0.0024, P=0.0040 for high-LG samples at P15 and low-LG samples at P21, respectively). *Ntrk2* mRNA levels showed statistically significant increases in ACX and VCX of low-LG and high-LG pups between P0 and P15, and between P0 and P21 (ordinary one-way ANOVA P=0.0004 and P=0.0023 for ACX and VCX, respectively; multiple comparisons P=0.0003, P=0.0056, for low-LG and high-LG samples in ACX, respectively; P=0.0056, P=0.0001, for low-LG and high-LG samples in VCX, respectively). It was noted that the relative levels of *Ntrk2* mRNA in the ACX were significantly different between low-LG and high-LG samples at P21 (boxed region in ACX *Ntrk2* panel of **Figure 7A**; multiple comparisons test P=0.0437).

**Figure 7B** shows developmental expression profiles for transcription factors *Fos*, *Jun*, *Nfkb1*, and *Otx2*. In the auditory brainstem the relative level of expression of *Fos* mRNA showed a small increase in CN of low-LG and high-LG pups between P0 and P21 (one-way ANOVA P=0.0032; and multiple comparisons test P=0.0065, P= 0.0021 for low-LG and high-LG samples, respectively). No significant changes in *Fos* mRNA were detected in pons between P0 and any other age examined (one-way ANOVA P=0.2129), while there was an increase in *Fos* mRNA in the IC of high-LG pups between P0 and P21 (one-way ANOVA P=0.0426; and multiple comparisons test P=0.0115). In contrast, *Fos* mRNA levels showed robust increases in ACX of pups from both LG groups between P0 and P15 (one-way ANOVA P=0.0148; and multiple comparisons test P=0.0060, P=0.0485, for low-LG and high-LG samples, respectively), and between P0 and P21 (multiple comparisons test P=0.0201, P=0.0360, for low-LG and high-LG samples, respectively). *Fos* mRNA levels increased in VCX of low-LG pups between P0 and P15 (one-way ANOVA P=0.0076; multiple comparisons P=0.0411), and in low-LG and high-LG pups between P0 and P21 (multiple comparisons test P=0.0029, P=0.0119, for low-LG and high-LG samples, respectively). In the auditory brainstem, the relative level of expression of *Jun* mRNA showed an increase in the CN and IC of low-LG and high-LG pups between P0 and P21 (one-way ANOVA P=0.0022, P=0.0002 for CN and IC, respectively; and multiple comparisons test P=0.0060, P=0.0002 for low-LG and high-LG CN samples, respectively; and P=0.0001, P=0.0012 for low-LG and high-LG IC samples, respectively). In the pons, *Jun* mRNA levels increased in low-LG pups between P0 and P15 (one-way ANOVA P=0.0001; and multiple comparisons test P=0.0246), and in both LG groups between P0 and P21 (multiple comparisons test P=0.0001, P=0.0001 for low-LG and high-LG pons samples, respectively). *Jun* mRNA levels showed significant increases in the ACX of pups from both LG groups between P0 and P15 (one-way ANOVA P=0.0001; and multiple comparisons test P=0.0106, P=0.0030, for low-LG and high-LG samples, respectively), and between P0 and P21 (multiple comparisons test P=0.0001, P=0.0001, for low-LG and high-LG samples, respectively). *Jun* mRNA levels increased in VCX of low-LG pups between P0 and P15 (one-way ANOVA P=0.0101; multiple comparisons P=0.0158), and in low-LG and high-LG pups between P0 and P21 (multiple comparisons test P=0.0032, P=0.0079, for low-LG and high-LG samples, respectively). The relative expression of *Nfkb1* mRNA did not change during development in the CN and pons (one-way ANOVA P=0.3487, P=0.3995, for CN and pons, respectively). In the IC, *Nfkb1* mRNA levels showed a slight but significant decrease in high-LG pups between P0 and P7 (one-way ANOVA P=0.0001; multiple comparisons test P=0.0100), and an increase in low-LG pups between P0 and P21 (multiple comparisons P=0.0001). The profile of *Nfkb1* mRNA expression was similar in the ACX and VCX, where there was an increase in low-LG pups between P0 and P15 (one-way ANOVA P=0.0001 and P=0.0001 for ACX and VCX, respectively; multiple comparisons test P=0.0157, P=0.0003 for low-LG samples in ACX or VCX, respectively), and an increase in pups from both LG groups between P0 and P21 (multiple comparisons test P=0.0001, P=0.0001, for low-LG and high-LG in ACX, respectively; and P=0.0001, P=0.0001, for low-LG and high-LG in VCX, respectively). It was noted that the relative levels of *Nfkb1* mRNA in the ACX were significantly different between low-LG and high-LG samples at P21 (boxed region in ACX *Nfkb1* panel of **Figure 6B**; multiple comparisons test P=0.0002). Lastly, the relative levels of expression of *Otx2* mRNA did not change during development in any of the brain structures examined (multiple comparisons test; P=0.4064, P=0.7841, P=0.4302, P=0.2075, P=0.4808, for CN, pons, IC, ACX and VCX, respectively).

**Figure 7C** shows mRNA developmental expression profiles for three downstream signaling effectors: the kinases *Akt1* and *Akt2*, and *Sort1*, a protein involved in the transport of other proteins from intracellular membrane compartments to the plasma membrane. In the CN, the relative levels of expression of *Akt1* increased between P0 and P15 in high-LG pups (one-way ANOVA P=0.0107; multiple comparisons test P=0.0003). It was noted that the relative level of *Akt1* mRNA was significantly different between low-LG and high-LG groups at P15 (boxed region in CN *Akt1* panel in **Figure 7C**; multiple comparisons test P=0.0030). In the CN, *Akt1* mRNA levels increased between P0 and P21 in both LG groups (multiple comparisons test P=0.0252, P=0.0101 for low=LG and high-LG samples, respectively). The levels of *Akt1* mRNA did not change during development in pons, IC, ACX and VCX (one-way ANOVA P=0.589, P=0.71611, P=0.2611 and P=0.1680 for pons, IC, ACX and VCX, respectively). In contrast to *Akt1* mRNA levels, the relative levels of *Akt2* mRNA increased in the CN, IC and VCX between P0 and P21 in both LG groups (one-way ANOVA P=0.0121, P=0.0012, P=0.0026 for CN, IC and VCX, respectively; multiple comparisons P=0.0020, P=0.0055 for low-LG and high-LG samples in CN; P=0.0004, P=0.0015, for low-LG and high-LG samples in IC; P=0.0457, P=0.0049, for low-LG and high-LG samples in VCX, respectively). In the pons, *Akt2* mRNA levels did not change between P0 and any age examined (one-way ANOVA P=0.6949). In the ACX, *Akt2* mRNA levels increased in high-LG pups between P0 and P15 (one-way ANOVA P=0.0001; multiple comparisons test P=0.0143), and in both LG groups between P0 and P21 (multiple comparisons test P=0.0001, P=0.0001, for low-LG and high-LG groups, respectively). Lastly, the levels of *Sort1* mRNA in the CN, ACX and VCX showed increased levels in both LG groups between P0 and P15 (one-way ANOVA P=0.0001, P=0.0001, P=0.0001, for CN, ACX and VCX, respectively; multiple comparisons test P=0.0321, P=0.0187 for low-LG and high-LG samples in CN, respectively; P=0.0050, P=0.0059 for low-LG and high-LG samples in ACX, respectively; and P=0.0017, P=0.0138, for low-LG and high-LG samples in VCX, respectively), and between P0 and P21 (multiple comparisons test P=0.0001, P=0.0001, for low-LG and high-LG samples in CN, respectively; P=0.0001, P=0.0001, for low-LG and high-LG samples in ACX, respectively; and P=0.0001, P=0.0001, for low-LG and high-LG samples in VCX, respectively). In the pons and IC, the levels of Sort1 mRNA increased in both LG groups between P0 and P21 (multiple comparisons test P=0.0001, P<0.0001, for low-LG and high-LG samples in pons, respectively; and P=0.0224, P=0.0002, for low and high-LG samples in IC, respectively). It was noted that mRNA levels in IC were statistically different between low-LG and high-LG samples at P21 (multiple comparisons test P=0.0329).

In sum, the data in **Figure 7** shows evidence that genes involved in transcriptional regulation and signal transduction the context of developmental plasticity, increased expression levels in the ACX and VCX between P0 and P21, and to some extent between P0 and P15. Similar age-dependent expression profiles were observed in the CN and IC, but not consistently in the pons. Statistically significant differences between samples from pups of low-LG and high-LG litters were observed in the ACX at P21 (*Ntrk2* and *Nfkb1*), in the CN at P15 (*Akt1*), and in the IC at P21 (*Sort1*).

### Analysis of genes involved in myelin development, the hypoxia-sensitive pathway and the Wnt7 pathway

**Figure 8** shows results for 10 genes that are involved in myelin development and two distinct signaling pathways. **Figure 8A** shows developmental expression profiles for *Olig2* and *Mbp*. In the CN, the relative levels of *Olig2* mRNA showed a significant increase in low-LG and high-LG samples between P0 and P21 (one-way ANOVA P=0.0250; multiple comparisons test P=0.0066, P=0.0067, for low-LG and high-LG samples, respectively), but there were no significant differences observed in the pons, IC, ACX and VCX between P0 and any age tested (one-way ANOVA, P=0.4959, P=0.7710, P=0.5170, P=0.3840, for pons, IC, ACX and VCX, respectively). The relative levels of *Mbp* mRNA showed an increase in the CN of low-LG pups between P0 and P21 (one-way ANOVA P=0.0068; multiple comparisons test P=0.0415). In the pons, *Mbp* mRNA levels decreased significantly in low-LG pups between P0 and P15 (one-way ANOVA P=0.3076; multiple comparisons test P=0.0238). In the IC, *Mbp* mRNA levels increased in high-LG pups between P0 and P21 (one-way ANOVA P=0.0073; multiple comparisons test P=0.0037). In ACX and VCX, *Mbp* mRNA levels increased in low-LG and high-LG pups between P0 and P21 (one-way ANOVA P=0.0002, P=0.0001 for ACX and VCX, respectively; multiple comparisons test P=0.0003, P=0.0004, for low-LG and high LG samples in ACX, respectively; P=0.0001, P=0.0002, for low-LG and high LG samples in VCX, respectively).

**Figure 8.**
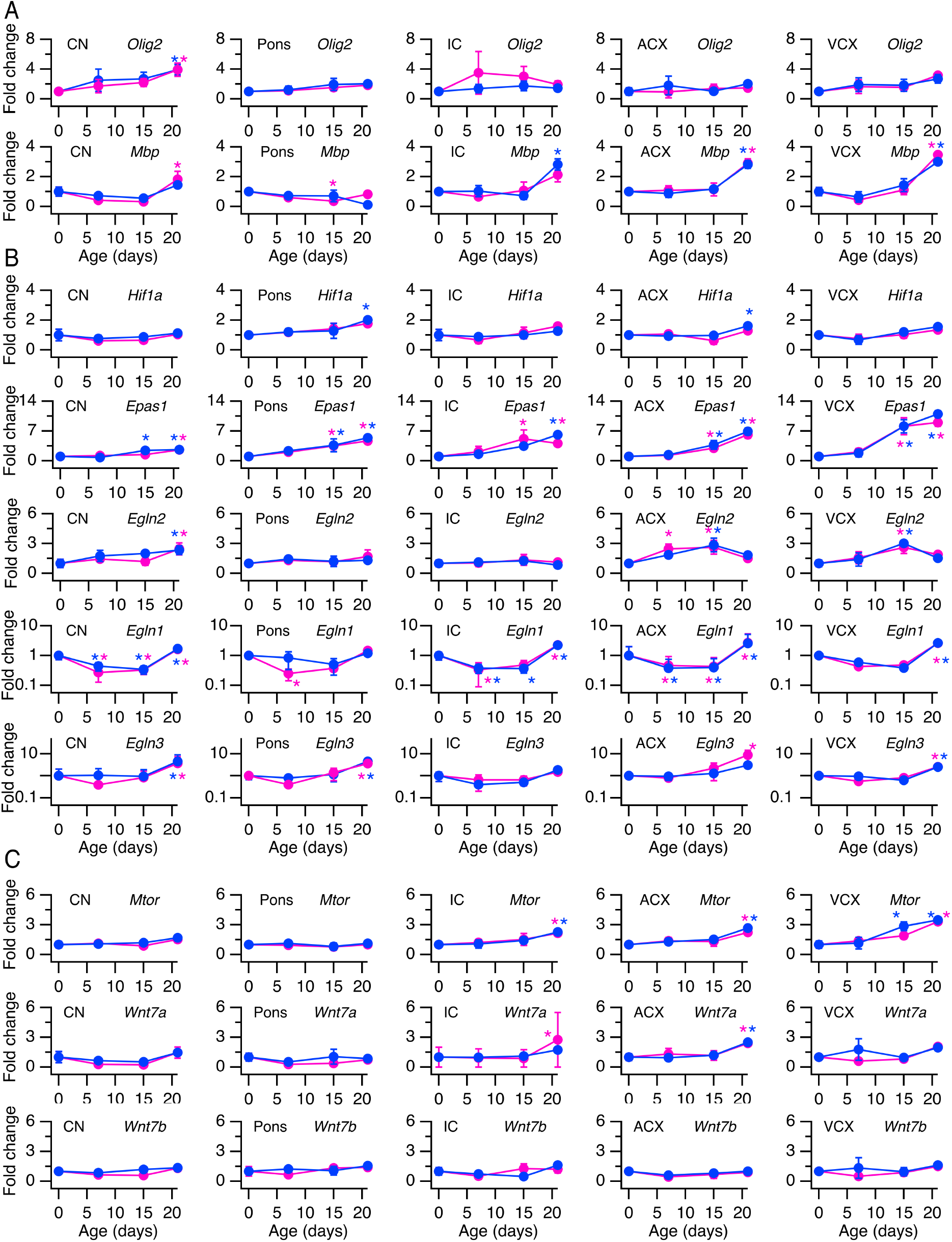
Temporal expression profiles for genes involved in oligodendrocyte development, the hypoxia-sensitive pathway and the mTor/Wnt7 pathway. **A,** Relative level of mRNA expression of Olig2 and Mbp. **B,** Relative level of mRNA expression of Hif1a, Hif2a, and Phd isoforms 1, 2 and 3. **C,** Relative level of mRNA expression of mTor, Wnt7a and Wnt7b. Magenta symbols represent data from low-licking and grooming (low-LG) samples. Blue symbols represent data from high-licking and grooming (high-LG) samples. Data represents mean ± sem (n=3 pups per age per LG group). Asterisks represent statistically significant differences with respect to P0. Alpha=0.05.

**Figure 8B** shows developmental expression profiles for hypoxia-sensitive transcription factor genes *Hif1a* and *Epas1* (*Hif2a*), and for *Egln* (*Phd*) paralogues 1-3 which code proteins that regulate Hif1a and Hif2a degradation. In the CN, IC and VCX, the relative levels of *Hif1a* mRNA did not change between P0 and any age examined (one-way ANOVA P=0.4775, P=0.3494, P=0.0716 for CN, IC and VCX, respectively). In the pons and ACX, a significant increase in *Hif1a* mRNA levels was detected in high-LG pups between P0 and P21 (one-way ANOVA P=0.1645, P=0.0699 for pons and ACX, respectively; multiple comparisons test P=0.0171, P=0.0203 for high-LG samples in pons and VCX, respectively). In the CN, pons, IC, ACX and VCX, the relative levels of *Epas1* mRNA showed increases between P0 and P15 and between P0 and P21 (one-way ANOVA P=0.0165, P=0.0303, P=0.0.0174, P=0.0001, P=0.0001 for CN, pons, IC, ACX and VCX, respectively). In the CN, an increase in *Epas1* mRNA in high-LG pups was observed between P0 and P15, while in the IC, an increase in low-LG pups between P0 and P15 was detected (multiple comparisons test P=0.0237, P=0.0092 for CN and IC, respectively). In the CN and IC, there was an increase of *Epas1* mRNA in both LG groups between P0 and P21 (multiple comparisons test P=0.0154, P=0.0119 for low-LG and high-LG samples in CN, respectively; P=0.0400, P=0.0021 for low-LG and high-LG samples in IC, respectively). In the pons, ACX, and VCX, an increase in *Epas1* mRNA was observed in both LG groups between P0 and P15 (multiple comparisons test P=0.0147, P=0.0105 for low-LG and high-LG pups in pons, respectively; P=0.0500, P=0.0104 for low-LG and high-LG pups in ACX, respectively; P=0.0006, P=0.0006 for low-LG and high-LG pups in VCX, respectively) and between P0 and P21 (multiple comparisons test P=0.0011, P=0.0003 for low-LG and high-LG pups in pons, respectively; P=0.0001, P=0.0001 for low-LG and high-LG pups in ACX, respectively; P=0.0002, P=0.0001 for low-LG and high-LG pups in VCX, respectively). The developmental expression profile of *Egln2* (*Phd1*) mRNA was heterogeneous. In the CN, there was a significant increase in low-LG and high-LG groups between P0 and P21 (one-way ANOVA P=0.1932; multiple comparisons test P=0.0303, P=0.0479 for low-LG and high-LG pups, respectively). In the pons and IC, there were no changes detected in *Egln2* mRNA levels between P0 and any age examined (one-way ANOVA P=0.9052, P=0.9349 for pons and IC, respectively). In the ACX, there were increases in *Egln2* mRNA of high-LG pups between P0 and P7 (one-way ANOVA P=0.0976, multiple comparisons test P=0.0351), and in both LG groups between P0 and P15 (multiple comparisons test P=0.0202, P=0.0096). In the VCX, there were increases in *Egln2* mRNA in both LG groups between P0 and P15 (one-way ANOVA P=0.0599; multiple comparisons test P=0.0205, P=0.0055). The developmental expression profile of *Egln1* (*Phd2*) mRNA levels was the first to show consistent changes between P0 and P7 across different brain regions. In the CN and ACX, there were decreases in *Egln1* mRNA levels in both LG groups between P0 and P7 (One-way ANOVA P=0.0001, P=0.0001; multiple comparisons test P=0.0050, P=0.0240, for low and high-LG samples in CN; P=0.0163, P=0.0066 for low-LG and high-LG samples in ACX), and in both LG groups between P0 and P15 (multiple comparisons test P=0.0084, P=0.0098 for low-LG and high-LG samples in CN; P=0.0119, P=0.0080 for low-LG and high-LG samples in ACX). In the CN and ACX, there were increases in *Egln1* mRNA levels in both LG groups between P0 and P21 (multiple comparisons test P=0.0156, P=0.0078 for low-LG and high-LG samples in CN; P=0.0001, P=0.0001 for low-LG and high-LG samples in ACX). In the pons, there was a decrease in *Egln1* mRNA levels in low-LG samples between P0 and P7 (one-way ANOVA P=0.0262; multiple comparisons test P=0.0452). In the IC, there were decreases in *Egln1* mRNA levels in both LG groups between P0 and P7 (one-way ANOVA P=0.0001; multiple comparisons test P=0.0267, P=0.0357), in high-LG pups between P0 and P15 (multiple comparisons P=0.0334), and increases in both LG groups between P0 and P21 (multiple comparisons P=0.0004, P=0.0005 for low-LG and high-LG samples respectively). Lastly, in the VCX there were increases in *Egln1* mRNA levels in both LG groups only between P0 and P21 (one-way ANOVA P=0.0001; multiple comparisons test P=0.0009, P=0.0007 for low-LG and high-LG samples, respectively). The developmental expression profile of *Egln3* (*Phd3*) mRNA in the CN, pons, and VCX, showed increases in both LG groups between P0 and P21 (one-way ANOVA P=0.0001, P=0.0001, P=0.0003 in CN, pons and VCX, respectively; multiple comparisons test P=0.0008, P=0.0.0001 for low-LG and high-LG samples in CN, respectively; P=0.0009, P=0.0001 for low-LG and high-LG samples in pons, respectively; P=0.0016, P=0.0020 for low-LG and high-LG samples in VCX, respectively). In the IC, there were no significant changes in *Egln3* mRNA between P0 and any age examined (multiple comparisons test all P values>0.05). In the ACX, there was an increase in *Egln3* mRNA levels between P0 and P21 in ACX of low-LG samples (multiple comparisons test P=0.0260).

**Figure 8C** shows developmental expression profiles for the kinase *Mtor*, and secreted signaling protein isoforms *Wnt7a* and Wnt7b. The relative levels of *Mtor* mRNA in the CN and pons did not change between P0 and any age tested (one-way ANOVA P=0.2815, P=0.7722 for CN and pons, respectively). In the IC and ACX, there were increases in both LG groups between P0 and P21 (one-way ANOVA P=0.1119, P=0.0056 for IC and ACX, respectively; multiple comparisons test P=0.0334, P=0.0210 for low-LG and high-LG groups in IC, respectively; P=0.0056, P=0.0005 for low-LG and high-LG groups in ACX, respectively). Lastly, in the VCX, there were increases in high-LG pups between P0 and P15 (one-way ANOVA P=0.0015; multiple comparisons test P=0.0057), and in both LG groups between P0 and P21 (multiple comparisons test P=0.0012, P=0.0007 for low-LG and high-LG samples, respectively). The developmental profile of *Wnt7a* mRNA in the CN, pons, and VCX, did not show any changes between P0 and any age examined (one-way ANOVA P=0.0682, P=0.5819, P=0.1640 for CN, pons and VCX, respectively). In the IC, there was an increase in *Wnt7a* mRNA in low-LG pups between P0 and P21 (one-way ANOVA P=0.0125; multiple comparisons test P=0.0023), and in the ACX, there were increases in *Wnt7a* mRNA in both LG groups between P0 and P21 (one-way ANOVA P=0.0153; multiple comparisons test P=0.0102, P=0.0059 for low-LG and high-LG samples, respectively). Lastly, there were no changes in *Wnt7b* mRNA levels between P0 and any age tested in any of the brain regions tested (one-way ANOVA P=0.1389, P=0.2215, P=0.2004, P=0.4577, P=0.6010 for CN, pons, IC, ACX and VCX, respectively).

In sum, the data in **Figure 8** shows evidence that genes involved in myelin development and two signaling pathways, the hypoxia-sensitive pathway and the *Mtor*/*Wnt7* pathway, *showed* heterogeneous changes, increasing between P0 and P15, between P0 and P21, or remaining constant throughout the ages examined. It is notable that mRNA levels for *Egln1* showed a decrease between P0 and P7, and between P0 and P15 in most auditory brain regions examined, but not in the VCX.

### Analysis of genes involved in cell signaling, water diffusion, and ion diffusion

**Figure 9** shows the developmental profiles of mRNAs coding for diverse membrane proteins involved in cell signaling, water, and ion diffusion. **Figure 8A** shows developmental profiles for genes of the vesicular glutamate transporter *Slc17a8* (*VGluT3*), the water channel *Aqp4*, the voltage-dependent potassium channel *Kcna3* (Kv1.3) and the voltage and calcium sensitive chloride channel *Ano1* (*TMEM16a*). The developmental profile of expression for *Slc17a8* mRNA did not show changes between P0 and any age tested in the CN, IC, ACX, and VCX (one-way ANOVA P=0.4894, P=0.4882, P=0.4543, P=0.7512 for CN, IC, ACX and VCX, respectively). In the pons, *Slc17a8* mRNA decreased between P0 and P15 in both LG groups (one-way ANOVA P=0.1447; multiple comparisons P=0.0126, P=0.0189 for low-LG and high-LG samples, respectively). The developmental profile of *Aqp4* mRNA expression in the CN showed increases between P0 and P7 in high-LG pups (one-way ANOVA P=0.0001; multiple comparisons test P=0.0402), between P0 and P15 in both LG groups (multiple comparisons test P=0.0012, P=0.0024 for low-LG and high-LG samples, respectively), and between P0 and P21 in both LG groups (multiple comparisons test P=0.0001, P=0.0001). In the pons, IC, ACX and VCX, there were consistent increases in *Aqp4* mRNA between P0 and P15 (one-way ANOVA P=0.0022, P=0.0010, P=0.0001, P=0.0001; multiple comparisons test P=0.0429, P=0.0248 for low-LG and high-LG samples in pons, respectively; multiple comparisons test P=0.0366, P=0.0079 for low-LG and high-LG samples in IC, respectively; multiple comparisons test P=0.0030, P=0.0037 for low-LG and high-LG samples in ACX, respectively; multiple comparisons test P=0.0143, P=0.0090 for low-LG and high-LG samples in VCX, respectively), and between P0 and P21 (multiple comparisons test P=0.0006, P=0.0008 for low-LG and high-LG samples in pons, respectively; multiple comparisons test P=0.0003, P=0.0003 for low-LG and high-LG samples in IC, respectively; multiple comparisons test P=0.0001, P=0.0001 for low-LG and high-LG samples in ACX, respectively; multiple comparisons test P=0.0001, P=0.0001 for low-LG and high-LG samples in VCX, respectively). The developmental expression profile for *Kcna3* mRNA in the CN, pons, IC, and VCX, did not show changes between P0 and any age tested (one-way ANOVA P=0.2693, P=0.6529, P=0.2559, P=0.3509 for CN, pons, IC and VCX, respectively). There was an increase in the ACX between P0 and P21 in high-LG samples (one-way ANOVA P=0.1153; multiple comparisons test P=0.0093). The developmental expression profile of *Ano1* mRNA in the CN, pons, IC, ACX, and VCX, did not show any changes between P0 and any age tested (one-way ANOVA P=0.0111, P=0.5035, P=0.6650, P=0.4616, P=0.3545 in CN, pons, IC, ACX and VCX, respectively).

**Figure 9.**
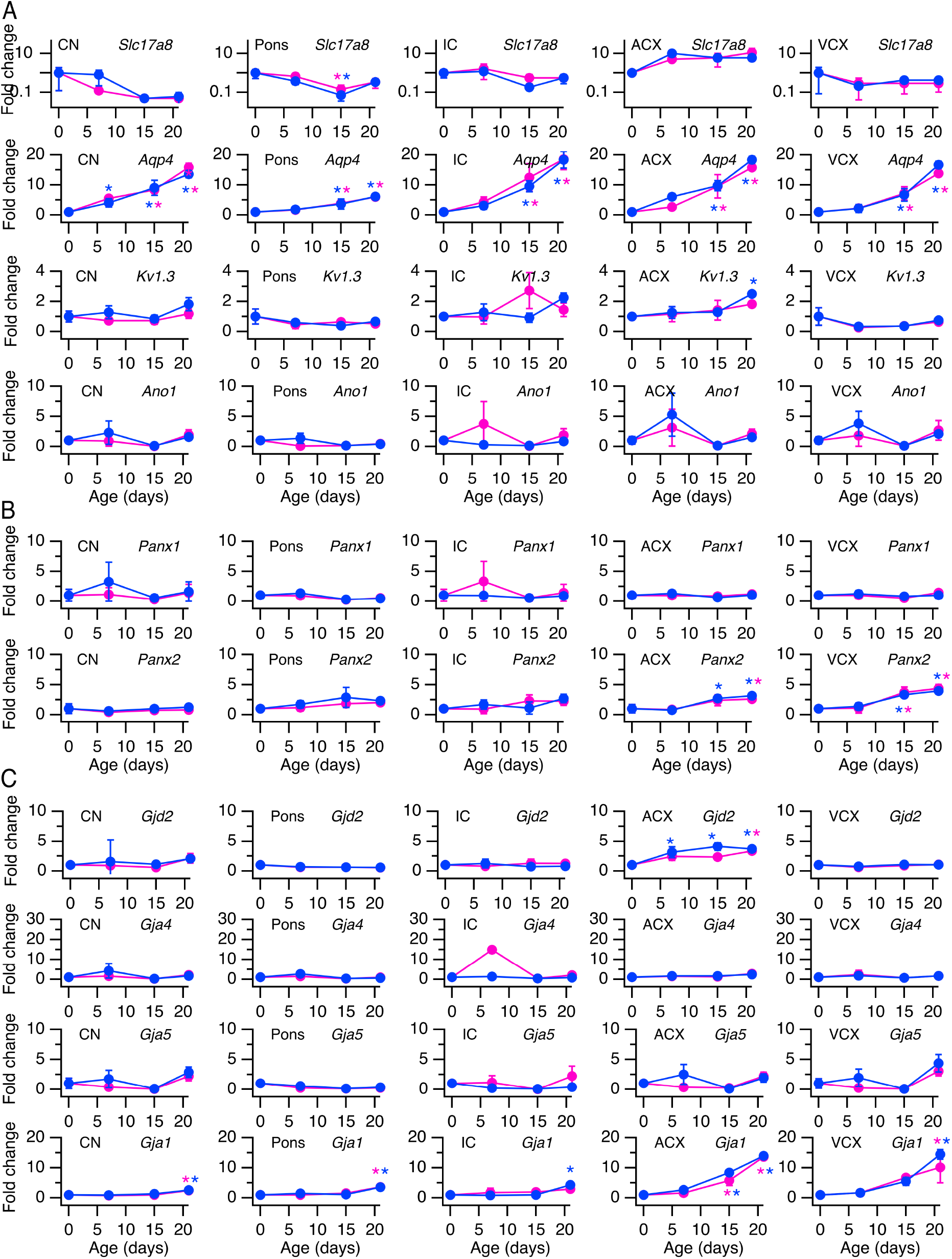
Temporal expression profiles for genes involved in neural signaling. **A,** Relative level of mRNA expression of Vglut3, Aqp4, Kv1.3, and TMEM16a. **B,** Relative level of mRNA expression of Panexin1 and Pannexin2. **C,** Relative level of mRNA expression of Cx36, Cx37, Cx40 and Cx43. Magenta symbols represent data from low-licking and grooming (low-LG) samples. Blue symbols represent data from high-licking and grooming (high-LG) samples. Data represents mean ± sem (n=3 pups per age per LG group). Asterisks represent statistically significant differences with respect to P0. Alpha=0.05.

**Figure 9B** shows the developmental expression profiles of *Panx1* and *Panx2* mRNAs. In the CN, pons, IC, ACX, and VCX, *Panx1* mRNA levels did not show any changes between P0 and any age tested (one-way ANOVA P=0.0237, P=0.1686, P=0.6542, P=0.8059, P=0.4887 for CN, pons, IC, ACX, and VCX, respectively). The developmental profile of *Panx2* mRNA did not show changes in auditory brainstem structures between P0 and any age tested (one-way ANOVA P=0.8168, P=0.5977, P=0.3431 for CN, pons, and IC, respectively). In contrast, in the ACX there were increases between P0 and P15 in high-LG pups (One way ANOVA P=0.0124; multiple comparisons test P=0.0272), and in VCX between P0 and P15 in both LG groups (one-way ANOVA P=0.0189; multiple comparisons test P=0.0226, P=0.0469 for low-LG and high-LG samples, respectively). Lastly, in the ACX and VCX, there were increases between P0 and P21 in both LG groups (multiple comparisons test P=0.0363, P=0.0076 in low-LG and high-LG groups in ACX, respectively; P=0.0072, P=0.0146 in low-LG and high-LG groups in VCX, respectively).

**Figure 9C** shows developmental expression profiles for the mRNAs of gap junction subunits *Gjd2* (*Cx36*), *Gja4* (*Cx37*), *Gja5* (*Cx40*) and *Gja1* (*Cx43*). In the CN, pons, IC, and VCX, the developmental profile of *Gjd2* expression did not show changes between P0 and any age tested (one-way ANOVA P=0.4181, P=0.6722, P=0.8983, P=0.9273 for CN, pons, IC and VCX, respectively). In contrast, there were increases in the ACX between P0 and P7 in high-LG pups (one-way ANOVA P=0.0202; multiple comparisons test P=0.0144), between P0 and P15 in high-LG pups (multiple comparisons test P=0.0012), and between P0 and P21 in both LG groups (multiple comparisons test P=0.0089, P=0.0032 for low-LG and high-LG samples, respectively). The developmental profile of expression of *Gja4* mRNA did not show any changes in the CN, pons, IC, ACX, and VCX, between P0 and any age tested (one-way ANOVA P=0.0245, P=0.3028, P=0.5237, P=0.5951, P=0.8205 for CN, pons, IC, ACX and VCX, respectively). Similarly, in the CN, pons, IC, ACX, and VCX, the expression levels of *Gja5* mRNA did not show any changes between P0 and any age tested (one-way ANOVA P=0.1392, P=0.4419, P=0.4332, P=0.1673, P=0.0261 for CN, pons, IC, ACX and VCX, respectively). In the CN, pons, and VCX, *Gja1* mRNA levels showed increases between P0 and P21 in both LG groups (one-way ANOVA P=0.0098, P=0.0001, P=0.0045; multiple comparisons test P=0.0145, P=0.0070 for low-LG and high-LG samples in CN, respectively; P=0.0001, P=0.0001 for low-LG and high-LG samples in pons, respectively; P=0.0099, P=0.0006 for low-LG and high-LG samples in VCX, respectively). In the IC, there was an increase between P0 and P21 in high-LG samples (one-way ANOVA P=0.0393; multiple comparisons test P=0.0058). Lastly, in the ACX, there were increases in both LG groups between P0 and P15 (one-way ANOVA P=0.0001; multiple comparisons test P=0.0012, P=0.0001 for low-LG and high-LG samples, respectively), and between P0 and P21 (multiple comparisons test P=0.0001, P=0.0001 for low-LG and high-LG samples, respectively).

In sum, the data in **Figure 9** shows evidence that the mRNA levels for several genes coding transmembrane proteins involved in cell signaling, water and ion diffusion did not change significantly between P0 and other ages examined. The exception was *Aqp4* mRNA, which showed consistent increased between P0 and P15, and between P0 and P21 in all brain regions examined.

## DISCUSSION

In this study we hypothesized that differences in maternal licking and grooming (LG) during the first week of life are associated with differences in the timing of hearing onset. However, the results of this study do not support the hypothesis, and indicate that different levels of maternal LG are not sufficient to modulate hearing onset in the progeny (**Figures 2 and 3**). Nevertheless, this study provides three new findings concerning auditory development: First, it shows that early onset of functional responses correlates with a variable delay to eye opening (EO; **Figure 4**); second, it adds new information on the relationship between the formation of the middle ear cavity and the threshold of functional responses during hearing onset (**Figures 5** and **6**); and third, it shows for the first time that mRNAs of the hypoxia-sensitive pathway and the *Bdnf* signaling pathway are regulated before and after the onset of hearing, respectively (**Figures 7–9**). Following is a discussion of the merits and limitations of these findings, including considerations for future studies.

### Variable delay between early hearing onset and EO

A major caveat of the approach used in the present study is that we were not able to predict the developmental profile of individual litters based on maternal LG. However, we were surprised to find that litters reared by low-LG and high-LG dams showed a similar range of early and late auditory brainstem response (ABR) onset. This turned out to be an advantage, because by tracking pups during development we were able to measure the delay between ABR onset and EO in a relatively large number of litters. The onset of hearing for airborne sounds precedes EO, and this sequence of events occurs prenatally or postnatally in different vertebrate species. The finding that litters with an early ABR onset have a more variable delay to EO (**Figure 4**) is relevant in the context of recent studies that manipulated the timing of EO and measured its effects on the development of membrane and synaptic properties of primary auditory cortex neurons in gerbils, and of synaptic properties of primary visual cortex neurons in Long-Evans rats (Mowery et al., 2016; Tatti et al, 2017). We propose that a better understanding of the relationship between hearing onset and the delay to EO will be useful to study cross modal experience-dependent plasticity between visual and auditory systems in rodents. Studies are needed to characterize the signaling pathways that influence the development of feed-forward and feedback mechanisms between ACX and VCX in animals with early and late hearing onset (Budinger et al., 2006; Mowery et al., 2016; Pan et al., 2018).

### Relationship between development of the auditory periphery and development of sensorineural responses from the inner ear

Combined functional and structural analyses from this study showed that cavity formation in the middle ear correlated with the type of sensorineural responses tracked in animals of different ages. We found that within a range of air volume from 15 mm^3^ to 40 mm^3^, ABRs had very elevated click intensity thresholds and relatively simple waveforms (**Figures 5 and 6**). For example, short latency potentials (SLPs) predominated over wave I responses at P12, and SLPs gradually waned as wave I responses increased in amplitude at P13 (**Figure 6**). Our results have confirmed and expanded on the previous findings of Blatchley, Cooper and Coleman, who described similar short latency responses to tone pips in ether-anesthetized P12 Sprague-Dawley rat pups (referred as summating potentials in their Figure 2: Blatchley, Cooper and Coleman, 1987). Additionally, it is puzzling to us that at P12 SLPs were observed without wave I responses, and that wave I responses were observed without subsequent ABR wave components II-V. Given the accumulating evidence that hair cells and neurons along the entire auditory system are functionally connected and active prior to hearing onset in different vertebrate species (Lippe, 1994; Gummer and Mark, 1994; Jones et al., 2007; Sonntag et al., 2009; Tritsch et al., 2010; Johnson et al., 2012; Babola et al., 2018; Corns et al., 2018), it is reasonable to propose that the functional changes observed in this study could reflect the contribution of conductive development to increased sensitivity (Saunders, Doan and Cohen, 1993), and the suppressive effects of isoflurane anesthesia on auditory function (Ruebhausen, Brozoski and Bauer, 2012; Bielefeld, 2014; Sheppard, Zhao, and Salvi, 2018). Future studies are needed to address how functional parameters of early sensory responses are affected by the state of the animal. This will require implementation of innovative methods to track structural and functional changes in non-anesthetized pups as they grow during postnatal development.

### Postnatal changes in gene expression of signaling pathways

The first postnatal weeks represent a sequence of sensitive periods when expression of genes involved in cellular proliferation, migration, differentiation, synaptogenesis, myelination, apoptosis, and neuroplasticity are regulated temporally and regionally in the CNS. The onset of ABRs and EO represent developmental stages in which auditory and visual experience can affect the above cellular processes in sensory pathways. In this study we found evidence that the hypoxia-sensitive pathway and the *Bdnf* pathway are regulated before and after the onset of hearing, respectively.

The hypoxia-sensitive pathway regulates gene expression by a negative feedback mechanism that is sensitive to a reduction in O_2_ partial pressure (i.e., hypoxia). O_2_ sensing is mediated by the products of *Egln* paralogue expression that interact with hypoxia-inducible factors coded by *Hif1a* and *Epas1* to target them for degradation in normoxic conditions. During hypoxic conditions Egln/Hif interactions decrease, reducing Hif degradation and increasing Hif stability that enables the transcriptional activation of the erythropoietin gene by Epas1, and the transcriptional regulation of genes encoding glycolytic enzymes by Hif1a (reviewed in Lappin and Lee, 2019). In this study we found that *Egln1* (*Phd2*) mRNA was down regulated in auditory brain regions before and after the onset of hearing and EO, while mRNAs for *Hif1a*, *Epas1*, *Egln2* and *Egln3* did not change, or increased at ages posterior to hearing onset (**Figure 8B**). This finding is intriguing to us and raises several questions for future studies: First, do cells that down regulate *Egln1* increase Hif activity? What causes down regulation of *Egln1* mRNA? What are the consequences of activating or disrupting the hypoxia-sensitive pathway during development of auditory brain regions? We propose that a first step to answer these questions will require mapping the localization of *Hif1a*/*Epas1* and *Egln1-3* mRNA and protein to specific cell types throughout the auditory system.

The Bdnf signaling pathway is involved in neuronal survival, cell growth, and differentiation via activation of its tyrosine kinase receptor Ntrk2, which in turn can modulate several signaling pathways including Akt/PI3K, Jak/STAT, NF-kB, UPAR/UPA, Wnt/β-catenin, and VEGF (Tajbakhsh et al., 2017). Studies in primary auditory cortex (ACX) and primary visual cortex (VCX) of rats and a mouse model of fragile X have shown that Bdnf signaling is regulated by sensory experience (Bozzi et al., 1995; Berardi, Pizzorusso and Maffei, 2000; Wang et al., 2017; Kulinich et al., 2019). In this study, we found a significant increase in mRNAs for *Ntrk2*, *Nfkb1*, *Akt2*, and *Wnt7a* in the ACX and VCX after hearing onset and EO (**Figures 7 and 8C**; although *Wnt7a* did not change in VCX). Furthermore, changes in mRNAs for *Ntrk2* and *Nfkb1* in the ACX were significantly different between pups reared by low-LG and high-LG dams at P21. These results expand previous findings that maternal care strongly modulates brain Bdnf levels in rodents (Branchi et al., 2013; Liu et al., 2000), and implicate maternal LG in experience-dependent development of functional responses in primary auditory cortex (de Villers-Sidani and Merzenich, 2011).

Lastly, in this study we found evidence of mRNA upregulation of alternative signaling pathways involving *Jun*, *Akt1*, and *Sort1* in subcortical auditory brain regions during a stage that coincides with the maturation of auditory thresholds and the end of the critical period for frequency tuning (Adise et al., 2014; de Villers-Sidani and Merzenich, 2011). Altogether, these results indicate a robust range of auditory periphery development and eye opening in Wistar rat pups that experience variation in maternal backgrounds. Consistent with this interpretation, there is a robust change in brain gene expression before hearing onset of *Egln1*, which codes for a crucial component of the hypoxia sensitive pathway. In addition, the results of this study implicate maternal LG in the expression of molecular factors involved in experience-dependent plasticity, neural signaling and transcriptional control in subcortical and cortical sensory brain regions of the progeny. Despite previous findings that maternal LG is increased in adoptive Wistar rat dams (Maccari et al., 1995), and that massage treatment during the sensitive period before EO accelerates development of visually evoked potentials in Long-Evans rats (Guzzeta et al., 2009), it is unlikely that increased physical stimulation through LG can explain the effects of maternal separation on ABR and middle ear development reported previously in Wistar rat pups (Adise et al., 2014).

## Acknowledgements

We would like to thank former lab members and students from Macaulay Honors Program Seminar 3 for discussions and help with behavioral scoring. Gene expression data was obtained and processed with help from the CUNY-ASRC Epigenetic Core facility staff. Supported by NIH grant SC1DC015907 and a CUNY ASRC-seed award.

